# Brain state shapes the polarity of pulsed infrared neural stimulation responses in individual cortical neurons

**DOI:** 10.64898/2026.09.19.752571

**Authors:** Peng Fu, Yin Liu, Liang Zhu, Mengqi Wang, Yuan Yu, Hequn Zhang, Yuchen Pei, Shy Shoham, Yan Zeng, Anna Wang Roe, Wang Xi

## Abstract

Pulsed infrared neural stimulation (INS, 1875nm the peak of energy absorption by water) is a non-viral method developed to activate single submillimeter sites in the brain. When used in ultrahigh-field fMRI, it reveals functionally specific brain-wide columnar networks comprising synaptically activated nodes. INS is non-damaging and effective for use in the human cortex and is useful for neural and behavioral neuromodulation in primates. To investigate how physiological state influences its cellular effects, we performed *in vivo* two-photon calcium imaging with GCaMP6s in mice and tracked identified cortical neurons in awake vs anesthetized states. Across the tested radiant exposures, INS-evoked responses differed markedly between states: under anesthesia, stimulation produced predominantly positive-going calcium responses, whereas during wakefulness the same identified neurons exhibited negative-going responses. Pharmacological manipulation of this baseline level by GABAergic signaling shifted response polarity in opposite directions: picrotoxin under anesthesia produced awake-like negative responses, whereas muscimol during wakefulness produced anesthetized-like positive responses. Trial-by-trial analyses further showed that the relationship between pre-stimulus baseline level and INS-evoked response depended on physiological state. Because INS relies on transient tissue heating, we also developed an *in situ* EGFP-based calibration approach to estimate temperature-related fluorescence changes. These fluoro-thermal contributions were modest at lower radiant exposures and became increasingly important at higher exposures, particularly near the fiber pathway. The low-intensity results are consistent with, but do not establish, a state-dependent feed-forward inhibitory framework. Together, these findings show that the neuronal response to focal INS is shaped by ongoing brain state and provide a practical framework for separating state-dependent calcium responses from temperature-related fluorescence effects, making INS a potentially versatile tool for neuroscience and clinical applications.

## Introduction

Neural stimulation can facilitate or interrupt the function of target brain areas, and thus potentially address specific clinical disorders by activating or disrupting neural activity^1–4^. Although both electrical and optical neural stimulation techniques have revolutionized experimental neuroscience and clinical neuromodulation^5–11^, they have shortcomings. Electrical stimulation has been the workhorse of brain stimulation, but is accompanied by current spread, which limits its spatial specificity and leads to potential side effects^5,6^. Modern optogenetic approaches offer exquisite cellular specificity, but are more challenging to implement in human tissue^9–11^. An alternative to these approaches is pulsed Infrared Neural Stimulation (INS, ∼1875 nm, peak of energy absorption by water), a promising opsin-free alternative with high spatial selectivity (submillimeter)^7,8^. INS is capable of evoking effective and non-damaging neuronal stimulation^12,13^, modulates both excitatory and inhibitory neurons^14^, and has been applied in multiple animal models (rodents^15,16^, cats^17^, primates^18,19^) as well as humans^13^. A primary distinction of INS studies from some other modalities (e.g., typical clinical electrical stimulation) is its submillimeter focality. Using specific pulsed parameters (0.2 ms pulse width, at 200 Hz for 0.5 s duration via a 200-μm diameter fiber optic), we have shown that INS selectively stimulates single cortical columns^17–21^ and submillimeter neuronal clusters in subcortical targets^22–24^. When INS is combined with optical imaging or ultrahigh-field MRI, systematic maps of local and brainwide columnar networks are revealed (e.g., reference^25^), thereby providing a view of the functionally specific brain circuits that mediate behavioral effects of focal stimulation. This approach of selecting functionally specific circuit-based stimulation sites introduces a promising route for effective therapy, one without the undesired side effects caused by recruitment of large, non-specific cell clusters and their respective circuits. However, precise spatial targeting does not necessarily guarantee predictable neural outcomes. A fundamental challenge in neuromodulation is understanding how stimulation parameters interact with the ongoing physiological state of the target neural circuit to shape neuronal responses.

One of the less understood aspects of brain stimulation is how physiological state influences stimulation outcomes. This issue is particularly important because many INS studies have been performed under anesthesia^14,16,18,26^, whereas clinical applications require predictable effects in awake subjects. Studies using electrical stimulation and transcranial magnetic stimulation have demonstrated that stimulation responses are shaped by the intrinsic state of neural circuits^27,28^. Electrical stimulation studies showed that elevated pre-stimulus activity can be associated with reduced evoked responses at the single-neuron level^27^, whereas TMS studies demonstrated that baseline cortical activity can predict stimulation efficacy^28^. These findings suggest that stimulation responses reflect the combined influence of external input and ongoing circuit state, raising the possibility that identical INS may generate different, or even opposite, neuronal responses under different physiological brain states (anesthesia vs. wakefulness). State-dependent modulation of INS has not been directly examined at the level of the same identified neurons. Moreover, previous work indicates that stimulation intensity influences the magnitude and polarity of INS-induced responses. For example, in the lower intensity range (<0.5 J/cm^2^), response amplitude correlates with increasing intensity, whereas at higher ranges, the effect can be reduced. Other infrared neural stimulation paradigms have also shown that the effect can be designed to be selective for fine fiber diameters^29^. A critical gap is therefore the lack of a direct within-neuron comparison of INS responses across awake and anesthetized states *in vivo*.

Here, we used *in vivo* two-photon calcium imaging to compare INS-evoked responses in the same cortical neurons across awake and anesthetized states. Previous studies from our group and others characterized INS-evoked calcium responses and their dependence on stimulation intensity, distance from the fiber tip, and neuronal population^14,26,30^. A second interpretational challenge arises from the thermal-mediated method of INS itself^31,32^, that is, transient heating can influence calcium fluorophores and thereby confound calcium measurements. We therefore addressed two linked questions: how physiological state shapes the polarity of INS-evoked neuronal calcium responses, and to what extent temperature-related fluorophore changes contribute to the measured signals. To address the latter, we developed an *in-situ* calibration framework for estimating fluoro-thermal fluorescence contributions. Resolving both issues is important for interpreting INS as a precise neuromodulation tool. Finally, we discuss a feed-forward inhibitory framework that is consistent with the present state-dependent data.

## Results

To characterize INS neuromodulation across physiological states, we performed two-photon calcium imaging of hSyn-GCaMP6s-labelled cortical neurons (mainly excitatory) in individual mice sequentially during awake and anesthetized conditions (**Fig. 1a-1c**). INS was first delivered in awake state using the established stimulation pattern (0.25 ms pulse width, 200 Hz, 0.5 s train duration, 2.5 s inter-train interval, radiant exposures 0.1-1.0 J/cm^2^ per pulse), which has previously been shown to modulate neurons at the stimulated site and at postsynaptic functionally connected sites^15,18,19,21,22^. This effective and non-damaging^12,13^ stimulation paradigm was also shown to induce neuronal calcium changes in anesthetized mice^14^. Specifically, as in our previous studies, using a 200-μm optical fiber targeting a focused area in the somatosensory cortex, INS trains were repeated 6 times, and calcium responses were imaged for 60 seconds per trial (**Fig. 1d**). This experimental design directly addresses a major limitation of previous INS studies, which were largely performed under anesthesia or relied on population-level measurements. By tracking identified neurons within the same cortical fields across awake and anesthetized states, calcium imaging revealed that response polarity is not an intrinsic property of stimulation, but instead emerges from the underlying circuit state, a distinction that cannot be readily resolved by conventional electrophysiological recordings.

**Fig. 1.**
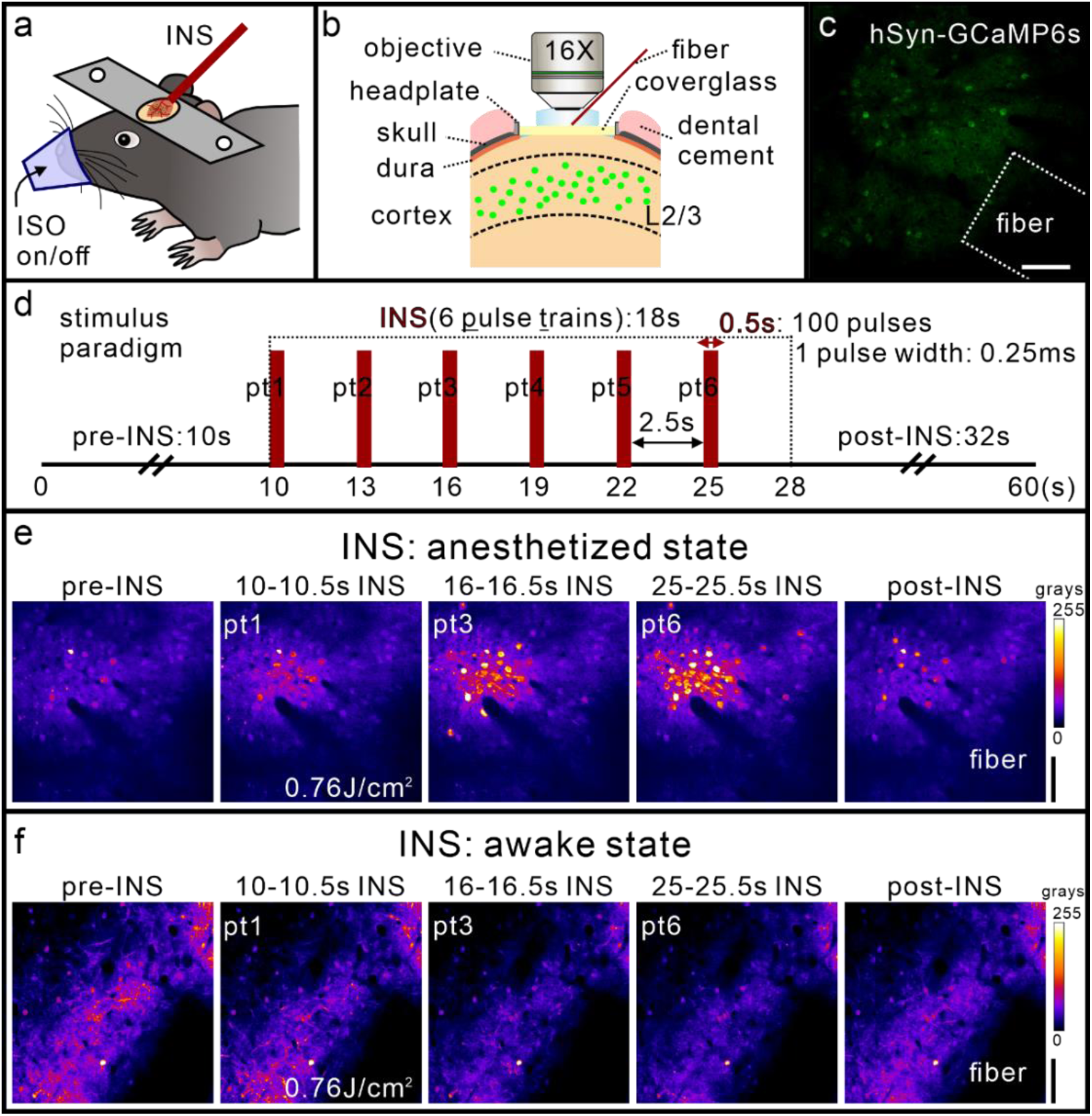
Two-photon calcium imaging reveals state-dependent neuronal responses to INS. **(a)** Imaging and stimulation setup under isoflurane (ISO) anesthesia. **(b)** Cranial window over mouse somatosensory cortex for two-photon imaging and infrared neural stimulation (INS). **(c)** hSyn-GCaMP6s-positive cortical neurons. Dotted outline: optical fiber tip. Scale bar, 100 μm. **(d)** INS stimulus paradigm. Each 60-s trial consisted of a 10-s pre-stimulation period, followed by six stimulation pulse trains (red bars, onset times below, delivered at 200 Hz for 0.5 sec, ISI 2.5 sec), and a 32-s post-stimulation recovery period. Seven radiant exposures were tested (0.16, 0.29, 0.42, 0.50, 0.59, 0.68, or 0.76 J/cm^2^) in pseudo-random order across trials. **(e)** Representative positive fluorescence responses to INS under anesthesia. Each image was averaged over the indicated time window. Scale bar, 100 μm. **(f)** Representative negative fluorescence responses to INS during wakefulness. Scale bar, 100 μm.

### Distinct calcium activity in response to INS during two states in the mouse cortex

In anesthetized mice, the averaged intensity map of neuronal response to each six-train INS stimulus (example shown in **Fig. 1e**, at the intensity of 0.76 J/cm^2^) produced a strong and focused calcium response (red-yellow in 10, 16, 25 s panels), after which neuronal calcium returned to its initial levels (post-INS). Activated neuronal clusters were about 200 μm in extent, consistent with our previous calcium imaging, intrinsic signal imaging, and functional MRI studies. In contrast, in the awake state, we observed a different scenario, with calcium activity decreasing during INS stimulation (example shown in **Fig. 1f**, at the intensity of 0.76 J/cm^2^, blue-purple region in 10, 16, 25 s panels. See time course at more intensities in **Supplementary Fig. 1a**) and gradually returning to baseline after stimulation.

To further examine this differential calcium activity, we compared the effects of INS on individual neurons in both awake and anesthetized states in the same field of view of the same mouse (**Fig. 2**). In awake mice, for a baseline reference image of the cortical region, we averaged spontaneous neuronal calcium activity over 60 seconds (**Fig. 2a**). In response to INS stimulation (**Fig. 2b**, 1 trial total 60 s: 10 s pre-INS baseline, 18 s INS period, 32 s post-INS period, at the intensity of 0.76 J/cm^2^), we found that neuronal calcium responses were modulated in phase with the pulse train delivery (each indicated by a dashed vertical line). Contrary to previous studies in anesthetized mice^14,16,26^, as shown in the four example neurons (1-4, labeled by white circles in **Fig. 2a**), there was a consistent *negative* response across the six successive pulse trains. Thus, the response of neurons in awake state appears to be a calcium signal with initial downward deflections from baseline, potentially indicating an ‘inhibition’ response^33,34^. These INS-induced negative calcium deflections in awake mice were confined to the focal region illuminated by the fiber tip and were not observed outside the illumination cone (**Supplementary Fig. 1b**, compare traces on left and right).

**Fig. 2.**
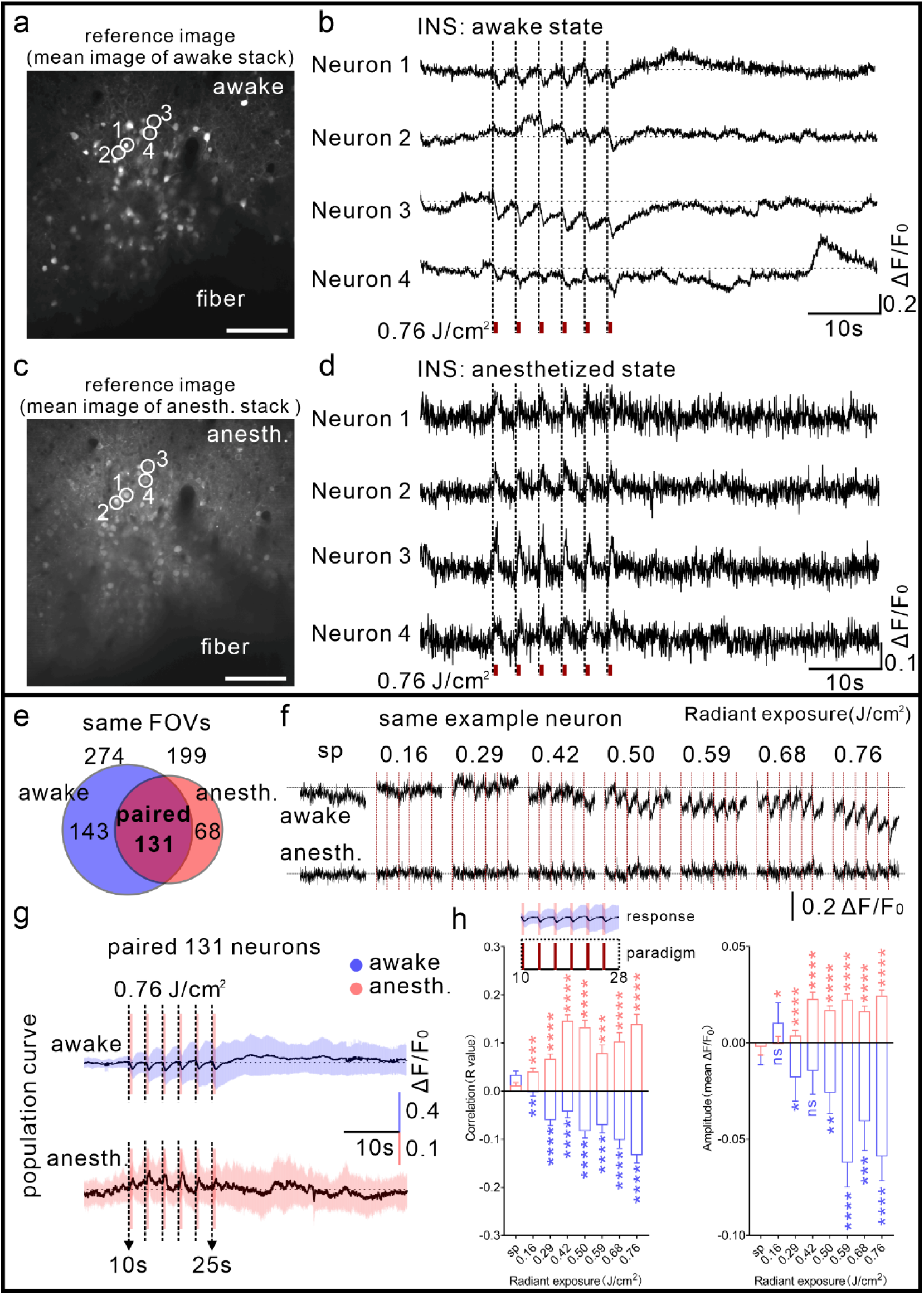
Distinct neuronal responses to INS in awake and anesthetized states of the same neurons. **(a)** Representative mean reference image stack during wakefulness (total six trials of image stack at intensity of 0.76 J/cm^2^). Scale bar, 100 μm. **(b)** Sample traces of negative calcium response to INS in awake state. Each trace was averaged from six trials for a single neuron. Four white circles indicated the selected neurons in (a). **(c)** Representative mean reference image stack under anesthesia (anesth.) after awake sessions in the same mouse (total six trials of image stack at intensity of 0.76 J/cm^2^). Scale bar, 100 μm. **(d)** Sample traces of positive calcium response to INS in anesthetized state. Four white circles indicated the selected neurons in (c), the same as in (a). **(e)** Paired neurons in the same field of view (n = 131 paired neurons from 3 mice) under awake and anesthetized states. Blue: awake, red: anesthetized, magenta: paired. **(f)** The example neuronal calcium activity in response to INS (during 10-28 seconds) across several intensities during awake and anesthetized states, respectively. The red vertical line indicates the INS period. **(g)** The example of population calcium activity of paired neurons (n = 131 paired neurons) in response to INS at the intensity of 0.76 J/cm^2^ under awake and anesthetized conditions. The black vertical line indicates the INS period. **(h)** Correlation and amplitude of responses to INS within population paired neurons across different intensity levels under awake and anesthetized states, respectively (paired Wilcoxon test between sp and INS). Data represent mean ± SEM in Supplementary Table 1. sp: spontaneous. anesth.: anesthetized.

We next examined responses of the same neuronal population after inducing isoflurane anesthesia in the same animals (**Fig. 2c**). The findings obtained under anesthesia were comparable to those of our earlier investigation^14^. That is, under anesthesia, in the same 4 neurons (1-4, labeled by white circles in **Fig. 2c**), INS induced a consistent positive response during the stimulation periods (also at the intensity of 0.76 J/cm^2^ in **Fig. 2d**). We compared the differences in spontaneous baseline activity in awake vs. anesthetized states. Awake mice had more active calcium activity in cortical neurons overall in spontaneous baseline activity, as well as more calcium events and longer calcium peak duration (**Supplementary Fig. 2a**, 274 neurons in awake state vs. 199 neurons in anesthetized state from three mice, see Methods for details). This reveals that mice in two states exhibit varying degrees of neuronal calcium activity, with awake mice exhibiting higher spontaneous calcium activity. We note that, in both awake and anesthetized states, INS stimulation did not alter spontaneous calcium activity in pre-vs. post-INS application, as evidenced by the number of calcium events and the duration of calcium peaks. (**Supplementary Fig. 2b and 2c** for awake and anesthetized states, respectively).

Below, we present population neurons’ calcium activity changes in both awake and anesthetized states in the same mouse. As described, we performed calcium imaging in three mice and recorded calcium activity from 274 neurons under awake conditions. Following anesthesia administration and repetition of the same experimental procedure within the identical field of views (FOVs), we detected 199 neurons. By aligning neuronal locations, we matched neurons between the awake and anesthetized states, resulting in a final set of 131 one-to-one matched neurons (**Fig. 2e**, magenta part). We present an example of a single, paired neuron recorded in awake and anesthetized states in turn (see top panel for awake state and bottom for anesthetized state in **Fig. 2f**). Its spontaneous activity and responses to INS stimulation at varying intensity levels are shown. The neuronal response to INS stimulation exhibited a progressive, intensity-dependent increase, with higher stimulation intensity eliciting stronger responses under both conditions.

At the population level, we extracted the calcium activity of these 131 paired neurons under each INS intensity level and computed their averages (as shown in **Fig. 2g** at the intensity of 0.76 J/cm^2^, and other INS intensities in **Supplementary Fig. 3a**). Consistent with the single-neuron observations described above, population neurons in awake animals exhibited a decrease in calcium activity during INS (blue curve), whereas these neurons in anesthetized animals showed an increase (red curve). To quantify the neuronal responses under both conditions, we analyzed the correlation and amplitude between neuronal calcium activity and the INS paradigm during the 10-28 second window (**Fig. 2h** top panel). The specific responses during INS were calculated in six INS pulse train periods (pt1, pt2, pt3, pt4, pt5, pt6 in **Supplementary Fig. 4**, more details in Methods), in which the R value of correlation coefficient was defined as correlation, and the averaged changes were defined as amplitudes, respectively. We found that under anesthesia, neurons exhibited a positive response (an increase in calcium signaling, see red bar chart), whereas in awake state, the response was negative (a decrease in calcium, see blue bar chart). In both states, however, the response magnitude strengthened in an intensity-dependent manner with increasing stimulation energy. The responses to INS between the two states were statistically significantly different (amplitude at more INS intensities in **Supplementary Fig. 3b**). Current evidence indicates that INS can modulate both excitatory and inhibitory neuronal populations^14^. However, the observed neural effects are strongly contingent upon the animal’s state (awake vs. anesthetized), likely reflecting net outcomes of state-dependent integrated circuit activity rather than direct, cell-type-specific actions. Taken together, these results demonstrate that INS exerts differential modulatory effects on neuronal calcium activity under different states. We next sought to quantitatively analyze the calcium activity of individual neurons across varying INS intensity levels in both awake and anesthetized conditions.

### Characterization of responses to INS across physiological states

Because individual neurons differed in the magnitude and direction of their awake-to-anesthetized response change, we used k-means clustering as a descriptive stratification of response patterns rather than as a definition of biological cell types (see example in single neuron in **Fig. 3a**). We found two main clusters of neurons based on the direction of change from awake to anesthetized state: 1) *Up neuron* (magenta dots) displayed responses with upwards change, and 2) *Other neuron* (grey dots) displayed minimal change. At 0.76 J/cm^2^, 53 of 131 paired neurons (40.4%) were assigned to an ‘Up neuron’ group exhibiting an upward change from awake to anesthetized state (**Fig. 3b** left, mean amplitudes changed from -0.181 ± 0.008 to 0.021 ± 0.005), whereas the remaining 78 neurons (59.6%) were grouped as ‘Other neuron’ and showed comparatively small mean changes (**Fig. 3b** right, mean amplitudes from 0.024 ± 0.014 to 0.027 ± 0.004). This suggests that INS has differential effects on individual neurons. Heatmaps of the paired neurons in a line-by-line manner illustrate the response pattern captured by this stratification in **Fig. 3c**. This finding indicates the consistency of the upward change of the group of Up neurons from awake to anesthetized states, interpreted as analysis-defined response groups.

**Fig. 3.**
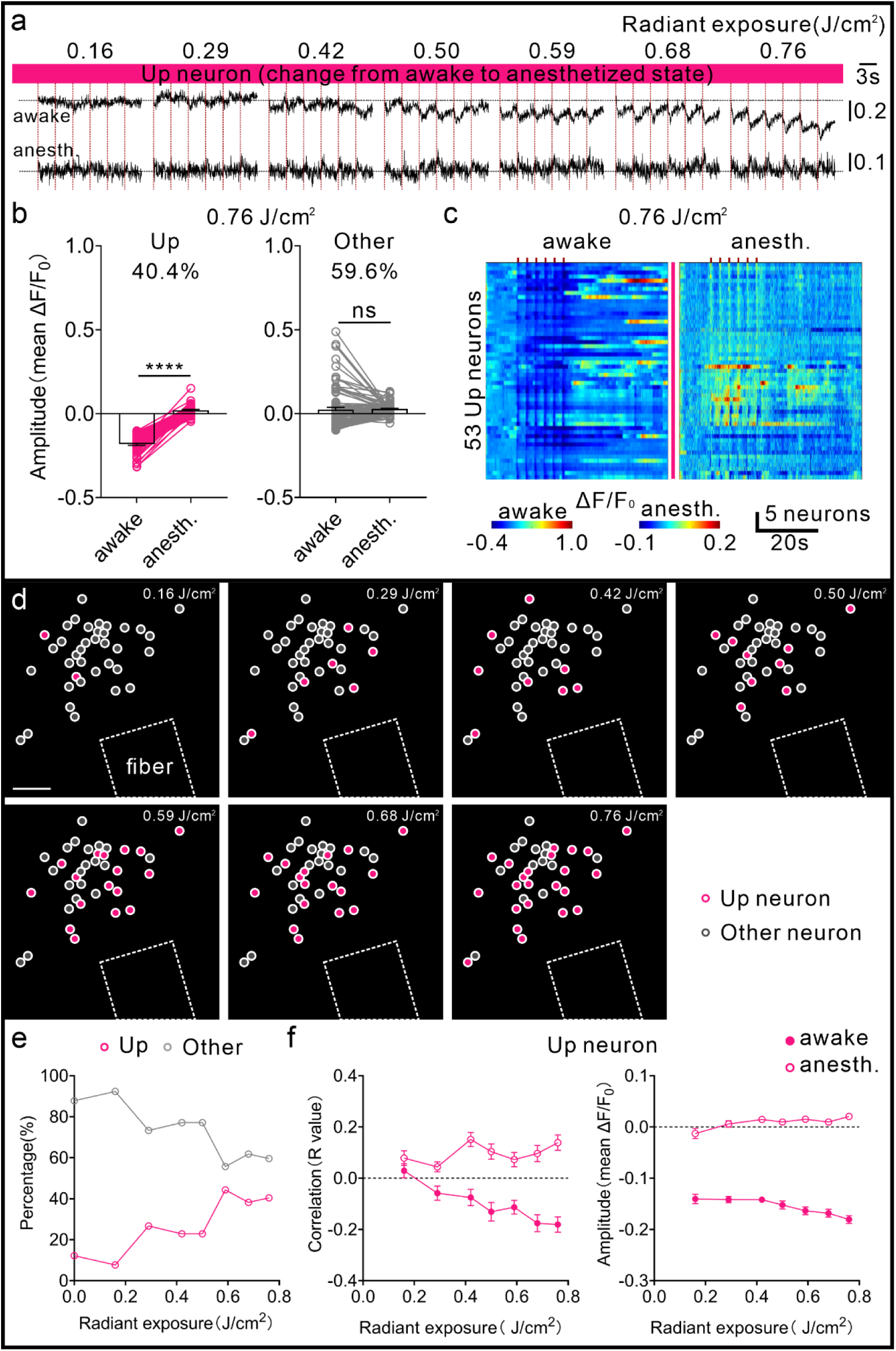
Descriptive response-pattern stratification of paired neurons across awake and anesthetized states. **(a)** The example timecourse (during 10-28 s) of an Up neuron across different intensities and states. The red vertical line indicates the INS period. **(b)** Category based on change direction of the amplitude of responses to INS from awake to anesthetized states (at the intensity of 0.76 J/cm^2^. Magenta, awake vs. anesth., paired Wilcoxon test, *p* < 0.0001; Gray: awake vs. anesth., paired Wilcoxon test, *p* = 0.0548). Magenta: Up neuron, gray: Other neuron. Data represent mean ± SEM. **(c)** All trial-averaged responses of paired neurons in mice (*n* = 53 Up neurons across three mice at the intensity of 0.76 J/cm^2^) and pseudo-colored heatmaps summarizing the calcium response trace for each neuron. The red bar indicates the INS period. **(d)** The categories of responses to different INS intensities between awake and anesthetized states in the distribution map of a single experimental animal. Dotted outline: optical fiber tip. Scale bar, 100 μm. **(e)** Proportion of each group across different INS intensity levels. **(f)** Correlation and amplitude of responses to INS within paired neurons in the Up groups across different intensity levels under awake and anesthetized states, respectively. Data represent mean ± SEM in Supplementary Table 2. anesth.: anesthetized.

We next examined how these operational response groups were distributed across the seven INS intensities (**Fig. 3d**). Both response patterns were spatially intermixed within the illuminated field of infrared light. The fraction of neurons assigned to the Up neuron generally increased with radiant exposure and was more stable across the higher intensities (0.59-0.76 J/cm^2^), whereas many neurons classified as Other neuron at lower intensities (0.16-0.50 J/cm^2^) were assigned to the Up group at higher intensities. This pattern suggests that the magnitude of the state-associated response shift is intensity-dependent, while avoiding interpretation of the clustering as evidence for fixed neuronal subtypes. At the population level, the proportion of neurons assigned to the Up group increased with stimulation intensity and approached approximately 40% at the higher exposures, while the proportion assigned to the Other group decreased (**Fig. 3e**). For the Up group, response correlation shifted from negative during wakefulness to positive under anesthesia, and response amplitude showed a corresponding upward shift across the tested intensities (**Fig. 3f**). These analyses provide a descriptive view of heterogeneity in state-associated INS responses.

Collectively, the paired-neuron data indicate substantial heterogeneity in how INS responses change between wakefulness and anesthesia in individual mice. This clear divergence not only underscores the state-dependent nature of INS modulation but also highlights its intensity-dependent regulatory effects, while reaffirming the reliability and reproducibility of INS-evoked calcium responses in neurons. To this end, we further focus subsequent mechanistic interpretation on the state-dependency of the hSyn-positive cortical population.

### State-dependent response shifts after pharmacological manipulation

Further experiments were designed to explore the mechanism underlying the state-dependent reversal in response polarity of excitatory neurons, which shifts from predominantly negative during wakefulness to positive under anesthesia. Specifically, we hypothesized that this divergence arises from differences in baseline neuronal activity, which is typically low under anesthesia and relatively high in the awake state. To test this, we conducted complementary pharmacological manipulations of GABAergic signaling in anesthetized and awake animals (**Supplementary Fig. 5a**). In the anesthetized condition, baseline activity was enhanced by local application of the GABA_A_ receptor antagonist picrotoxin (PTX, 100 μM, see **Fig. 4**) through the coverglass with a hole within the cranial window (**Supplementary Fig. 5b-5c** for setup). Conversely, in awake animals, baseline activity was suppressed by application of the GABA_A_ receptor agonist muscimol (MSM, 100 μM, see **Fig. 5**).

**Fig. 4.**
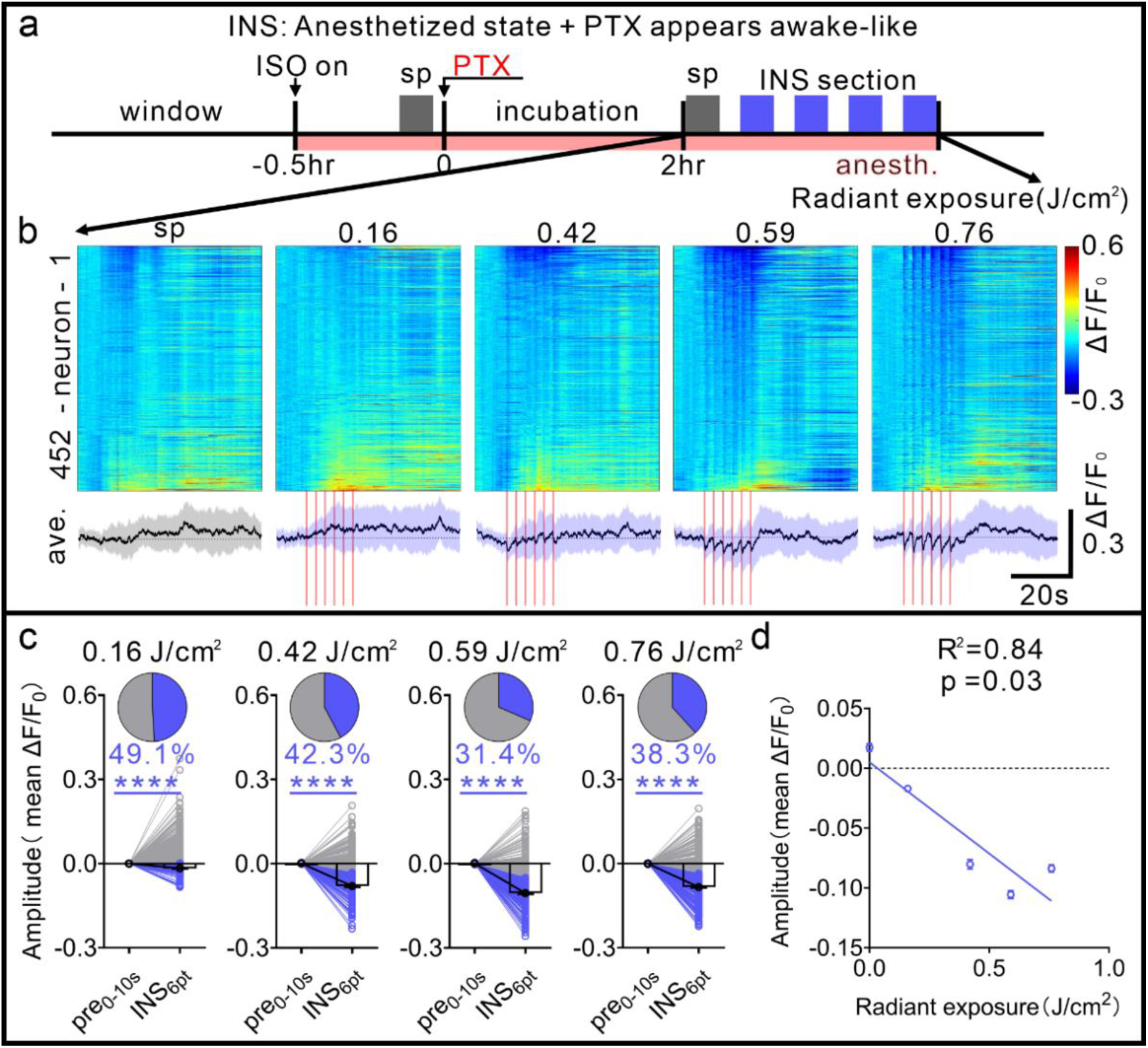
PTX-associated shift of INS responses toward an awake-like pattern under isoflurane anesthesia. **(a)** Pharmacological picrotoxin (PTX) manipulation timeline in anesthetized mice. **(b)** Top panel, all trial-averaged responses of neurons in PTX-manipulated mice (*n* = 452 neurons across three mice) and pseudo-colored heatmaps summarizing the calcium response trace for each neuron across spontaneous and four INS-induced activities. Bottom panel, a grand average of spontaneous (gray) and INS-induced (blue) calcium activity for all neurons in awake-like state. The pink bar indicates the INS period. The shadow indicates SD. **(c)** The paired mean responses of significantly reversed neurons (blue line) during the INS-period showed a negative deflection in anesthetized state with PTX across different intensities. Black line, averaged response of the pre-INS baseline and INS stage (blue, pre vs. INS, paired Wilcoxon test, *p* < 0.0001). **(d)** The correlation between laser intensity and the magnitude of the negative transient neural responses. (Y = -0.094×X + 0.0249, *p* = 0.0326). Data represent mean ± SEM in Supplementary Tables 3 and linear regression in Supplementary Table 4. sp: spontaneous; PTX: picrotoxin; anesth.: anesthetized.

**Fig. 5.**
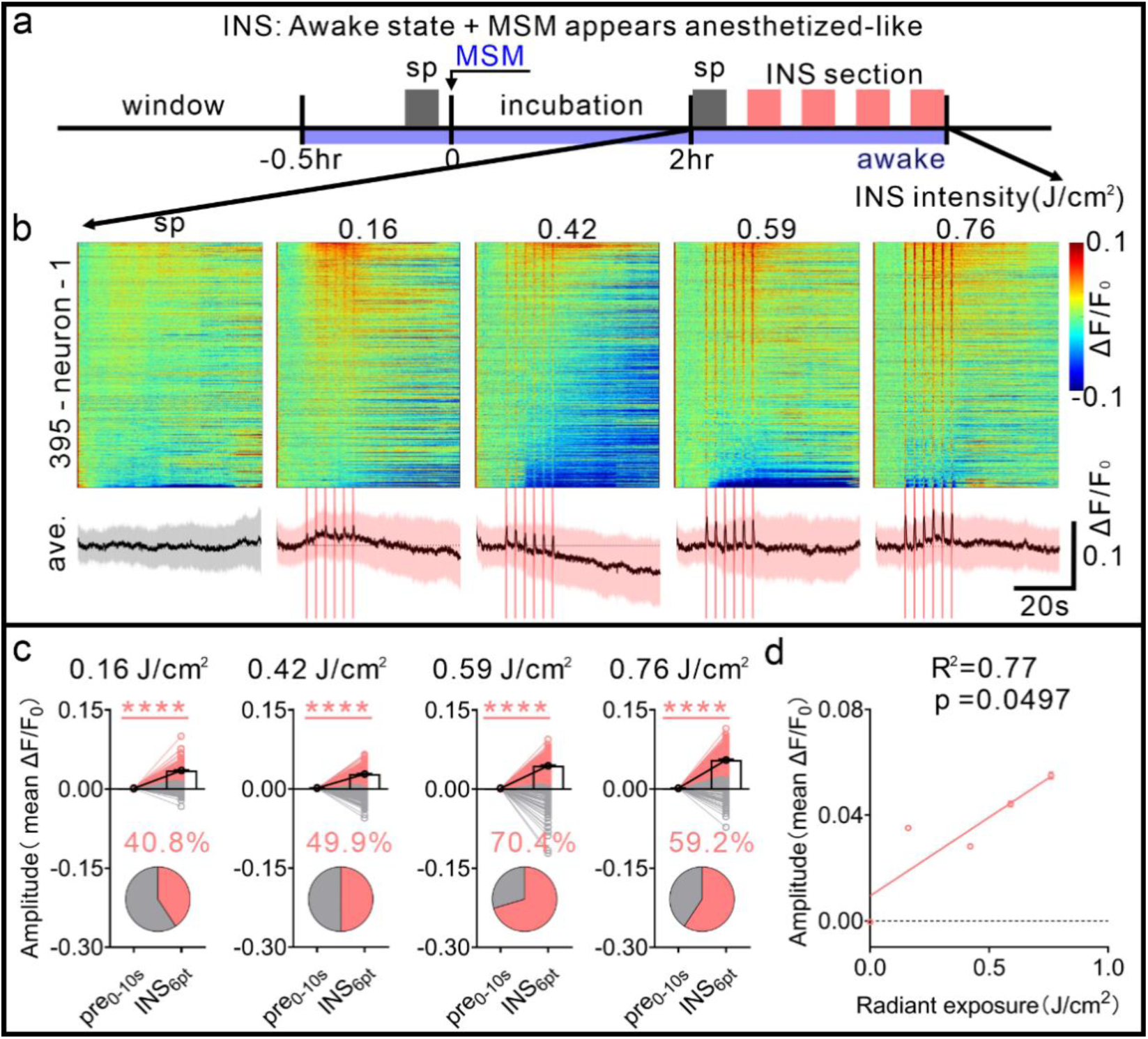
MSM-associated shift of INS responses toward an anesthetized-like pattern during wakefulness. **(a)** Pharmacological muscimol (MSM) manipulation timeline in awake mice. (**b)** Top panel, all trial-averaged responses of neurons in MSM-manipulated mice (*n* = 395 neurons across three mice) and pseudo-colored heatmaps summarizing the calcium response trace for each neuron across spontaneous and four INS-induced activities. Bottom panel, a grand average of spontaneous (gray) and INS-induced (red) calcium activity for all neurons in the MSM-manipulated anesthetized-like state. The pink bar indicates the INS period. The shadow indicates SD. **(c)** The paired mean responses of significantly reversed neurons (red line) during the INS-period showed a positive deflection in awake state with MSM across different intensities. Black line, averaged response of the pre-INS baseline and INS stage (red, pre vs. INS, paired Wilcoxon test, *p* < 0.0001). **(d)** The correlation between laser intensity and the magnitude of the positive transient neural responses. (Y = 0.039×X + 0.0031, *p* = 0.0494). Data represent mean ± SEM in Supplementary Tables 5 and linear regression in Supplementary Table 6. sp: spontaneous; MSM: muscimol.

As a timeline of application in **Fig. 4a**, we first utilized PTX in anesthetized mice (maintained with 0.6% isoflurane mixed with fresh air) to increase the level of baseline neuronal activity, thereby mimicking an ‘awake-like’ state^35,36^. Following PTX application, spontaneous baseline activity (sp, left column) and INS-induced activity (INS, four intensities displayed in second to fifth columns as 0.16, 0.42, 0.59, and 0.76 J/cm^2^) were measured in **Fig. 4b** (*n* = 452 neurons from 3 mice). Application of PTX led to an increase in spontaneous activity compared with anesthetized state (see first column and **Supplementary Fig. 6a-6b**). Importantly, in contrast to our results from the isoflurane-controlled anesthetized state (see Fig. 2g red curves in population data), INS stimulation resulted in a *decreased* response of calcium activity in anesthetized mice with PTX incubation (see second to fifth columns), similar to the response of neurons in awake state (see Fig. 2g blue curves in population data). Based on the amplitude calculation of the k-means clustering, shifted INS-evoked responses toward negative deflections resembling the pattern observed during wakefulness (**Fig. 4c** blue part). As illustrated in Fig. 4c after clustering, the amplitude during the six INS pulse trains that exhibited decreased response was similar to that in awake state (blue bar, compare with Fig. 2h blue bar). The magnitude of the negative response generally increased across the tested radiant exposures in anesthetized mice after PTX application, although the relationship was not strictly monotonic at the highest intensity (**Fig. 4d**, linear fit, R^2^ = 0.84). This evidence supports that the role of GABA_A_ in the decreased baseline activity present under anesthesia was associated with an awake-like shift in INS response polarity.

**Fig. 6.**
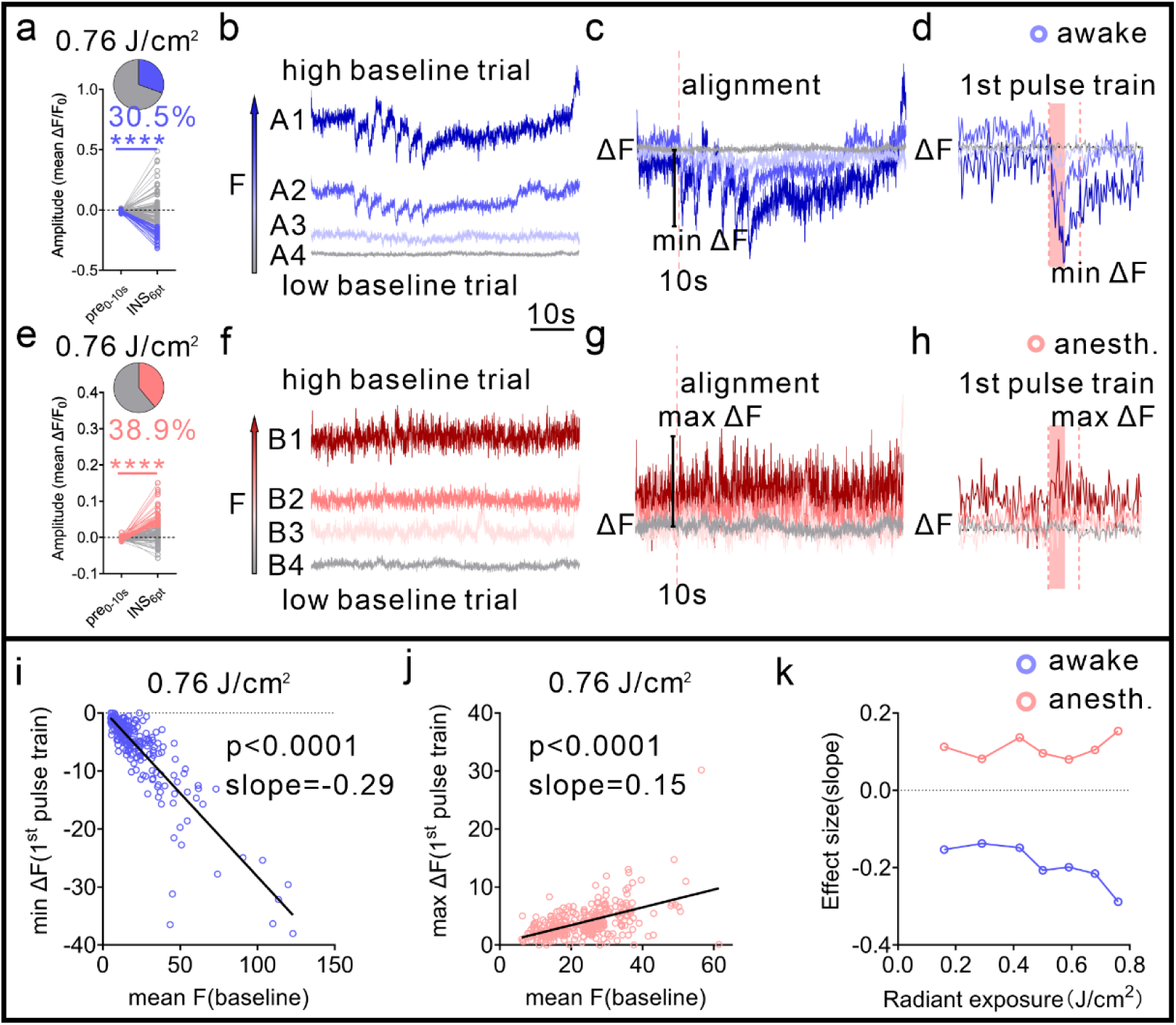
Pre-stimulus baseline is associated with INS-evoked responses in a state-dependent manner. **(a)** Example response-clustered neuronal group at 0.76 J/cm^2^ showing negative INS-evoked deflection in awake state (pre vs. INS, paired Wilcoxon test, *p* < 0.0001). **(b-c)** Example trial-by-trial relationship between raw pre-stimulus baseline (F) and subsequent fluorescence change (ΔF) in awake state. **(d)** The calculation diagram of the association between the response of the pre-stimulus baseline level (0-10 s) and the first stimulation epoch (10-11 s, INS delivered from 10-10.5 s) for the awake state. **(e)** Example response-clustered neuronal group at 0.76 J/cm^2^ showing positive INS-evoked deflection in anesthetized state (pre vs. INS, paired Wilcoxon test, *p* < 0.0001). **(f-g)** Corresponding examples under anesthesia. **(h)** The calculation diagram of the baseline-response association for the anesthetized state. **(i)** Negative association in the selected awake-response group at 0.76 J/cm^2^ (240 trials from 40 neurons across three mice, Y = -0.29×X + 0.61, *p* < 0.0001). **(j)** Positive association in the selected anesthetized-response group at 0.76 J/cm^2^ (306 trials from 51 neurons across the same mice, Y = 0.15×X + 0.35, *p* < 0.0001). **(k)** Regression-slope effect-size index across stimulation intensities for awake (blue) and anesthetized (red) states. anesth.: anesthetized. Data represent mean ± SEM in Supplementary Table 7 and linear regression in Supplementary Table 8.

In the complementary experiment, to mimic the ‘anesthetized-like’ state, we examined the response to MSM in awake mice to decrease the baseline level of neuronal activity^37–39^. After MSM incubation in awake animals (timeline as shown in **Fig. 5a**), the spontaneous activity (sp, left column, lower level of spontaneity, see **Supplementary Fig. 6c-6d**) and INS-induced activity (INS, four intensities shown in second to fifth columns) were recorded. We found that the neurons in awake state with MSM (*n* = 395 neurons from 3 mice) exhibited a *positive* calcium response (**Fig. 5b**, second to fifth columns), which showed a response similar to neurons in anesthetized state (see Fig. 2h red bar). In MSM-manipulated awake mice, many neurons showed increased calcium response to INS and a few neurons showed decreased response, a result similar to that in actual anesthetized mice. Moreover, INS-evoked responses shifted toward positive deflections resembling those recorded under anesthesia (**Fig. 5c** red part). Consistent with our results from anesthetized animals, the increase in calcium response was consistent with the increase in INS intensities in awake mice after MSM application (shown in **Fig. 5d**, linear fit, R^2^ = 0.77). Taken together, both PTX (awake-like) and MSM (anesthetized-like) experiments show that bidirectional state manipulations of the baseline level through GABA_A_ receptor signaling can shift INS response polarity in opposite directions. Because these interventions simultaneously alter inhibitory tone, spontaneous baseline activity, and network excitability, the results support a state-dependent circuit interpretation but do not establish baseline calcium activity as the sole mediator.

### Pre-stimulus activity is associated with INS responses in a state-dependent manner

We next asked whether trial-to-trial variation in the pre-stimulus baseline level was associated with the response to the first INS train. To minimize potential carry-over from preceding pulse trains, we related the pre-stimulus period (0-10 s) to the first stimulation epoch (10-11 s, INS delivered from 10-10.5 s) in awake and anesthetized recordings. This analysis was intended to characterize a state-dependent association rather than to establish pre-stimulus baseline level as a direct measure of membrane potential or as a causal determinant.

For visualization and regression analyses, we examined response-defined groups identified by the clustering procedure: neurons with negative INS responses during wakefulness and neurons with positive responses under anesthesia (**Fig. 6a** for awake state, **Fig. 6e** for anesthetized state). Trial-by-trial examples illustrate how raw pre-stimulus fluorescence (F) and the subsequent fluorescence change (ΔF) covaried within individual neurons. In awake recordings, an example response trial with four different pre-stimulus baseline levels (F, 4 trials of A1-A4 in **Fig. 6b**) was plotted, in which the dark blue trial represents a high baseline level and the light blue trial represents a low baseline level. We then aligned the response curve to the time point just before INS stimulation (at 10 s, pink dotted vertical line) as the subsequent fluorescence change (ΔF, see **Fig. 6c**), and calculated the change in negative response in the first stimulation epoch (min ΔF in awake state). As shown in **Fig. 6d**, higher pre-stimulus baseline was associated with larger negative deflections in the selected response group. Similarly, we performed the same operation on anesthesia data. In anesthetized recordings, higher pre-stimulus baseline was associated with larger positive deflections (4 trials of B1-B4 in **Fig. 6f-6h**, max ΔF in anesthetized state). Due to raw fluorescence can also reflect indicator expression, optical factors, and recording stability, we interpret these analyses as within-dataset associations with pre-stimulus baseline rather than as direct measurements of neuronal membrane potential.

To summarize these relationships across intensities, we used the slope of the trial-by-trial linear regression between the pre-stimulus baseline level and the first-train response as an effect-size index (**Fig. 6i-6k**). Negative slopes were observed for the selected awake-response group, whereas positive slopes were observed for the selected anesthetized-response group (**Supplementary Fig. 7** and **Fig. 8**). At 0.76 J/cm^2^, the corresponding slopes were approximately -0.29 and 0.15, respectively (240 trials in awake state and 306 trials in anesthetized state). These analyses indicate that the sign of the association between pre-stimulus baseline level and INS-evoked response differs between the two physiological states (**Fig. 6k**, blue curve for awake state and red curve for anesthetized state). Beyond the biological state-dependent modulation of INS-evoked responses, an important methodological question is to estimate the fluoro-thermal fluorescence in pulsed INS application.

### In situ calibration of INS-induced fluoro-thermal fluorescence changes

Given the state-dependent reversal of GCaMP6s responses, we next quantified the fluoro-thermal fluorescence component expected during pulsed infrared stimulation. We imaged cortical neurons expressing EGFP in awake mice and used this calcium-insensitive GFP-family reporter to estimate temperature-dependent fluorescence dynamics *in situ*^40^ (**Fig. 7a**). Across four INS intensities, EGFP fluorescence showed time-locked negative deflections that increased with pulse energy (**Fig. 7b**, *n* = 803 hSyn-EGFP-positive neurons from 3 mice), consistent with the known negative temperature dependence of GFP fluorescence (approximately -0.9%/°C^41^, **Fig. 7c**). The spatial distribution of the EGFP fluorescence dips was intensity-dependent. At the intensity of 0.42 J/cm^2^, fluorescence decreases were modest across the field of view, indicating little localized thermal contribution (**Fig. 7d**). At the intensity of 0.76 J/cm^2^, dips were spatially heterogeneous and largest near the fiber pathway, with some cells reaching approximately 13% fluorescence decrease, consistent with a focal thermal gradient (**Fig. 7e**).

**Fig. 7.**
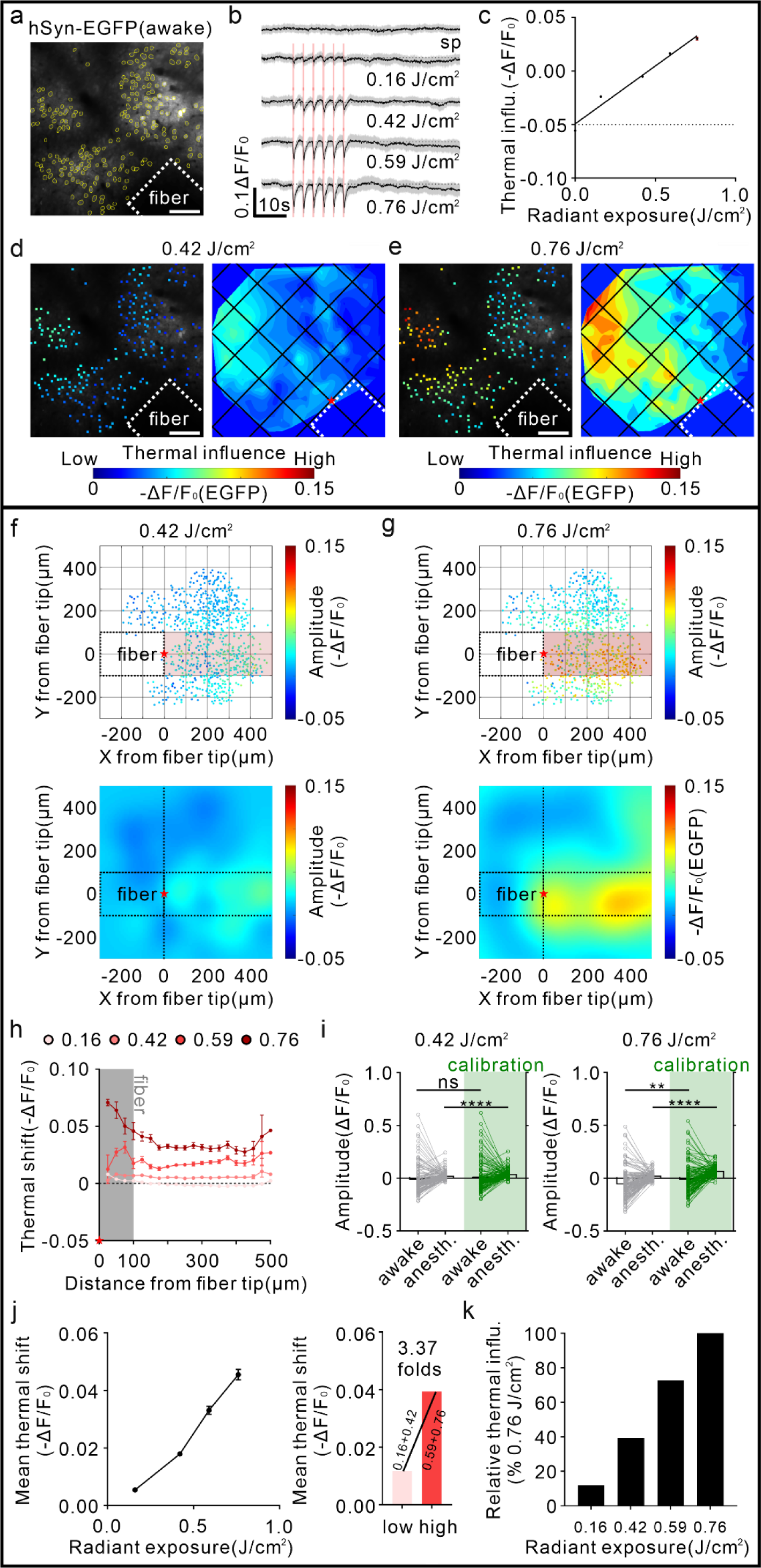
In situ calibration of INS-induced fluoro-thermal contributions in cortical neurons. **(a)** Experimental schematic for calibrating the thermal-induced fluorescence changes using EGFP-expressing neurons in awake mouse somatosensory cortex. The white dotted box marks the position of the INS fiber tip. Neuronal somas were segmented using Cellpose. Scale bar, 100 μm. **(b)** Averaged population time traces of EGFP fluorescence dips synchronized with pulsed INS delivery across four increasing intensities in awake mice (n = 803 neurons from 3 animals). Time-series data are presented as mean ± SD. **(c)** The negative fluorescence changes scale monotonically with INS intensity, consistent with the established temperature-dependent quenching coefficient of EGFP fluorescence (Y = 0.108×X + 0.0006, R^2^ = 0.97, p = 0.0018). **(d-e)** Single-cell fluorescence change maps at representative low (0.42 J/cm^2^, d) and high (0.76 J/cm^2^, e) intensities. Scale bar, 100 μm. (**f-g**) Gaussian-smoothed spatial maps of EGFP fluorescence change after pooling single-neuron coordinates from all three animals and normalizing positions relative to the fiber tip origin (marked by a red star in the maps), shown for 0.42 J/cm^2^ (f) and 0.76 J/cm^2^ (g), respectively. These maps illustrate the radial spread of the thermal artifact around the stimulation pathway (pink shaded bars). (**h**) Distance-dependent EGFP fluorescence-change profiles across 0.16-0.76 J/cm^2^, plotted as a function of radial distance from the fiber tip. (**i**) EGFP-based estimate of fluoro-thermal contribution to GCaMP6s recordings (0.42 J/cm^2^: -0.015 ± 0.012 (awake) vs. 0.003 ± 0.012 (awake calibration), Mann-Whitney test, *p* = 0.0688; 0.023 ± 0.003 (anesth.) vs. 0.041 ± 0.003 (anesth. calibration), Mann-Whitney test, *p* < 0.0001; 0.76 J/cm^2^: -0.059 ± 0.013 (awake) vs. -0.014 ± 0.012 (awake calibration), Mann-Whitney test, *p* = 0.0029; 0.025 ± 0.003 (anesth.) vs. 0.070 ± 0.004 (anesth. calibration), Mann-Whitney test, *p* < 0.0001), used to distinguish thermal fluorescence quenching from state-dependent neuronal calcium responses. (**j**) EGFP-based thermal shift across INS radiant exposures. The mean thermal shift at the two higher radiant exposures (0.59 and 0.76 J/cm^2^) was approximately 3.37-fold greater than that at the two lower radiant exposures (0.16 and 0.42 J/cm^2^). (**k**) Relative thermal influence (RTI) across INS radiant exposures. RTI was calculated by normalizing the EGFP-based thermal shift at each radiant exposure to the maximal thermal shift measured at 0.76 J/cm^2^, which was defined as 100%. The relative thermal influence increased from 11.9% and 39.3% at 0.16 and 0.42 J/cm^2^ to 72.7% and 100% at 0.59 and 0.76 J/cm^2^, respectively. Data represent mean ± SEM. anesth.: anesthetized.

We then pooled single-neuron coordinates from all three animals, normalized positions relative to the fiber tip, and generated Gaussian-smoothed spatial maps of EGFP fluorescence change at representative lower and higher intensities (0.42 and 0.76 J/cm^2^, **Fig. 7f** and **7g**, respectively). Distance-dependent profiles across the tested intensity range (0.16-0.76 J/cm^2^) showed that the thermal fluorescence component was most prominent within approximately 100 μm of the fiber tip at higher intensities (>0.5 J/cm^2^) and decreased rapidly with distance (**Fig. 7h**).

Finally, we used the EGFP measurements as an empirical estimate of the direction and magnitude of possible fluoro-thermal contributions to the GCaMP6s recordings (**Fig. 7i**). The magnitude of the EGFP-based thermal correction increased progressively with radiant exposure, from 0.0054 ±0.0003 at 0.16 J/cm^2^ to 0.0455 ±0.0018 at 0.76 J/cm^2^ (**Fig. 7j**). When normalized to the maximal thermal shift at 0.76 J/cm^2^, the relative thermal influence increased from 11.9% and 39.3% at 0.16 and 0.42 J/cm^2^ to 72.7% and 100% at 0.59 and 0.76 J/cm^2^, respectively (**Fig. 7k**). Accordingly, the mean thermal shift at the two higher radiant exposures was approximately 3.37-fold greater than that at the two lower exposures. Note that in practice, in both anesthetized^17,21^ and awake^42^ states, the range of intensities used is 0.1-0.3 J/cm^2^, values that lead to heat increases not exceeding 0.5 ℃ as shown by MR thermometry^43^. The comparison indicates that lower-intensity awake-state negative GCaMP6s responses are unlikely to be explained by temperature-dependent fluorescence changes alone, whereas interpretation of higher-intensity responses should account for an increasing focal thermal component. Moreover, the same EGFP-derived correction was applied to awake and anesthetized datasets, the estimated thermal shift was independent of physiological state, whereas the underlying GCaMP6s response polarity remained state-dependent. Because EGFP and GCaMP6s measurements were obtained from separate cohorts, this calibration provides a population-level estimate rather than a direct per-cell correction. It therefore defines a practical framework for assessing when neuronal calcium responses are likely to dominate over fluorophore-related thermal effects.

### Conceptual model of neuronal calcium responses to INS in distinct brain states

To place the state-dependent responses in a circuit framework, we considered a minimal feed-forward inhibition (FFI) model in which a common excitatory input recruits both a pyramidal neuron and an inhibitory interneuron, with the interneuron providing feed-forward inhibition to the pyramidal neuron^44^. This model is presented as a conceptual framework rather than as a directly measured circuit in the present experiments of INS-induced responses. In particular, the hSyn-GCaMP6s recordings do not identify a specific excitatory-cell subtype, and membrane potential was not measured directly.

We utilized this FFI model in an attempt to understand the state-dependent effects during INS experiments. Moreover, in view of previous analogous studies of brain stimulation, we compared the neuronal calcium responses to INS to the responses to high-frequency electrical stimulation (ES)^27^. In such an FFI framework, the balance between excitatory and inhibitory synaptic components can vary with the operating state of the postsynaptic neuron^45,46^, and these measured synaptic responses are relative adaptations while altering the potential levels of the neuron (V_m_) over a wide range^47^. With these, when at relatively depolarized states (V_m_ = -70 mV), excitatory postsynaptic potentials (EPSPs) dominate the evoked postsynaptic potentials and inhibitory postsynaptic potentials (IPSPs) are minimized after ES, causing burst firing in pyramidal neurons (**Fig. 8a**). Whease at the depolarized level (e.g., V_m_ = -50 mV, see **Fig. 8c**), ES to the tissue evokes a compound synaptic response that consists of a short EPSP followed by a long-lasting IPSP (can be contributed by the activation of GABA_A_ and sometimes GABA_B_ receptors). We therefore use the schematic in Fig. 8 to illustrate one possible way in which the same external input could produce different net calcium responses to INS across physiological states. The negative responses observed during wakefulness (**Fig. 8b**) and positive responses observed under anesthesia (**Fig. 8d**) are consistent with this general framework, but the schematic membrane-potential levels and synaptic components are not direct measurements from the present study. Direct electrophysiological measurements of membrane potential and synaptic excitation/inhibition will be required to test this mechanism.

**Fig. 8.**
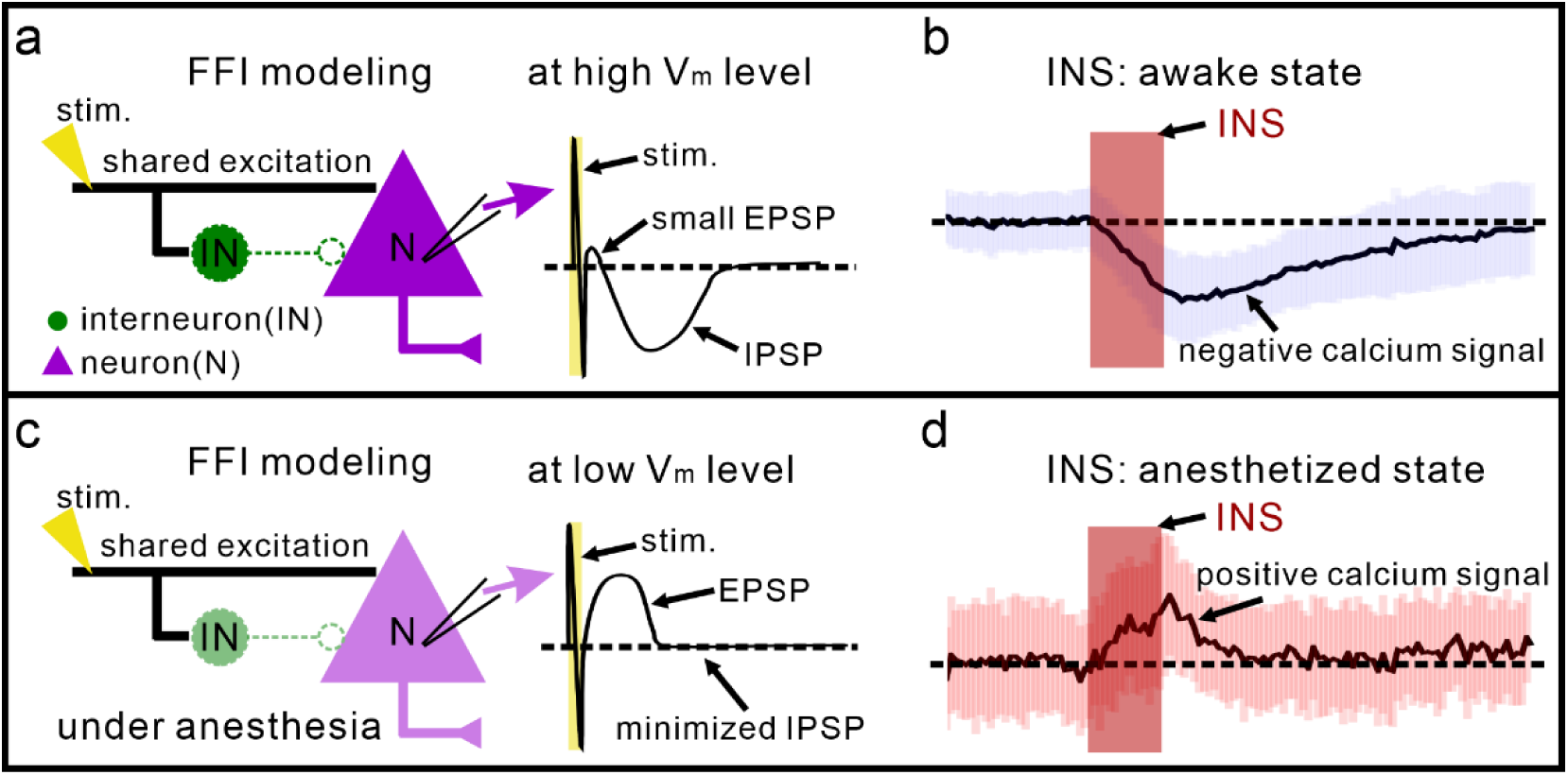
Conceptual framework for state-dependent INS responses. **(a)** Schematic minimal feed-forward inhibition (FFI) circuit in a relatively depolarized operating state (high level, -50 mV). A shared excitatory projection (black) drives an interneuron (green) and a pyramidal neuron (purple), with feed-forward inhibition from the interneuron to the pyramidal neuron. The accompanying EPSP/IPSP trace of the compound synaptic response to electrical stimulation (ES) is schematic and illustrates a condition in which inhibitory influence is relatively prominent. **(b)** Representative negative GCaMP6s fluorescence response to INS during wakefulness. **(c)** The same conceptual circuit in a relatively hyperpolarized operating state (low level, -70 mV), with a schematic synaptic response to ES in which the EPSP component is relatively more prominent. **(d)** Representative positive GCaMP6s fluorescence response to INS under anesthesia. stim.: stimulation; EPSP: excitatory postsynaptic potential; IPSP: inhibitory postsynaptic potential.

## Discussion

### Summary

In this study, combining two-photon calcium imaging with pharmacological manipulations of GABAergic signaling, we show that INS-evoked calcium responses can reverse in polarity at the single-cell level between anesthetized and awake states. Tracking identified neurons within the same cortical fields showed positive calcium deflections under anesthesia and negative deflections during wakefulness under otherwise matched stimulation conditions. With further pharmacological manipulation of brain state, GABA_A_ receptor blockade with picrotoxin under anesthesia shifted responses toward the awake-like negative pattern, whereas GABA_A_ receptor activation with muscimol during wakefulness shifted responses toward the anesthetized-like positive pattern. These pharmacological effects, together with the trial-by-trial associations with pre-stimulus fluorescence, support a role for ongoing circuit state but do not establish baseline calcium activity as a sole determinant. A second contribution of this study is an *in situ* EGFP-based calibration framework showing that fluoro-thermal effects become increasingly important at higher radiant exposures. Together, the data indicate that INS responses are shaped by physiological state and that thermal fluorescence contributions should be explicitly considered when interpreting optical readouts. A feed-forward inhibitory circuit provides one plausible conceptual framework for these observations.

### State-dependent inversion of cortical responses to brain stimulation

Neuromodulatory responses to electrical stimulation (ES) and transcranial magnetic stimulation (TMS) have been shown to depend strongly on the physiological state of the brain^27,28,48–52^. However, these observations are difficult to compare directly because ES and TMS recruit neuronal populations over substantially different spatial scales and through different biophysical mechanisms, ranging from relatively local axonal activation to widespread network modulation. Consequently, whether state-dependent effects reflect intrinsic circuit properties or simply differences in the populations recruited by each stimulation modality remains unresolved. For example, two-photon imaging during ES in awake mice demonstrated that pre-stimulus activity and cell-type-specific recruitment shape stimulation outcomes, with elevated baseline activity associated with reduced evoked responses in both excitatory and inhibitory neurons^27^. Conversely, TMS studies in anesthetized cats showed that higher pre-stimulus activity predicted stronger post-stimulation responses^28^. Rather than indicating a single universal relationship between baseline activity and stimulation efficacy, these findings suggest that the effect of ongoing activity depends on both the physiological state of the circuit and the stimulation context.

Our study extends this framework to INS by showing that the same identified neurons can exhibit opposite response polarities across awake and anesthetized states while stimulation location and physical parameters remain unchanged. The paired-neuron design is important because it reduces variation arising from neuronal identity, recording location, and stimulation geometry, allowing physiological state to be evaluated within the same local circuit. The observed polarity reversal was associated with pre-stimulus baseline activity and could be shifted in opposite directions by pharmacological manipulation of GABAergic signaling. Thus, the response to INS appears not to be an invariant consequence of infrared energy delivery, but rather an outcome shaped by the interaction between the external stimulus and the ongoing state of the cortical circuit.

Several alternative explanations and design limitations should be considered. First, negative fluorescence deflections during wakefulness could arise partly from temperature-dependent changes in the fluorescent reporters rather than neuronal calcium dynamics. The *in situ* EGFP measurements demonstrate that such fluoro-thermal effects are present, especially at higher radiant exposures, but are insufficient to explain the lower-intensity state-dependent polarity reversal on their own. Second, awake recordings preceded anesthetized recordings in the paired experiments, so physiological state is not completely separable from session order. The pseudo-randomized stimulation intensities, recovery periods, within-neuron tracking, and pharmacological response shifts reduce but do not eliminate this potential confound. Third, the principal datasets contain many neurons and repeated trials nested within a relatively small number of animals; consequently, neuron- and trial-level statistics should be interpreted in the context of biological replication across mice. Larger cohorts, counterbalanced state order, and hierarchical statistical models will be important for future confirmation. Finally, anesthesia alters multiple aspects of cortical physiology, including membrane potential, synaptic transmission, and excitation-inhibition balance. The present results therefore support state-dependent circuit processing but should not be interpreted as identifying a single baseline variable pathway as the sole cause of response polarity.

### Thermal contributions and inhibitory mechanisms of INS

A particular interpretational challenge for INS is that its biophysical action depends on transient tissue heating, while fluorescence from GFP-family reporters is itself temperature sensitive^41,53^. A decrease in GCaMP fluorescence during INS therefore cannot, by itself, be interpreted as neuronal inhibition. As a second important contribution of this study, we developed an *in situ* calibration framework for quantifying temperature-related fluorescence changes during INS. To address this issue directly, we used EGFP as a calcium-insensitive reporter to estimate the fluorescence component associated with heating under the same stimulation conditions. The resulting signal was both intensity- and distance-dependent: temperature-related fluorescence decreases were modest at lower intensities (roughly 15% at 0.2 J/cm^2^ and 40% at 0.4 J/cm^2^), but became increasingly prominent near the optical fiber at higher intensities (roughly 70% at 0.6 J/cm^2^ and 100% at 0.8 J/cm^2^). This spatial and intensity dependence is consistent with a localized thermal contribution and defines an important limitation for interpreting calcium imaging during high-intensity INS. Importantly, however, the magnitude of the EGFP fluorescent changes at lower intensities was insufficient to reproduce the negative responses observed in awake GCaMP6s-expressing neurons. Thus, the fluorescence decrease measured during wakefulness cannot be attributed solely to direct temperature dependence of the calcium reporter. Combined with the fact that increasing INS intensities in anesthetized state leads to increasing positive response, the idea that INS effects are primarily thermal is not supported. We also note that, in practice, in both anesthetized^17,21^ and awake^42^ states, the range of intensities used is 0.1-0.3 J/cm^2^, values that fall within the range of modest heat effect (this study) and lead to heat increases not exceeding 0.5 ℃ as shown by MR thermometry and Monte Carlo simulations^21,43^.

This distinction is important when considering previous studies of inhibitory effects of infrared stimulation. INS has been reported to enhance inhibitory synaptic activity^54^, suppress fine axonal excitability^29^, and reduce neural firing^55^ in several preparations. These findings demonstrate that infrared stimulation is capable of engaging inhibitory processes, but they do not imply that inhibition is a universal or direct consequence of INS. The reported effects vary substantially across species, preparations, fiber geometry, stimulation parameters, and physiological states. For example, suppression of firing has also been reported in anesthetized rat cortex using a substantially larger optical fiber, where broader recruitment and surround inhibition may contribute to the observed response^15,18^. Such variability is therefore more consistent with a context-dependent balance between excitation and inhibition than with a fixed inhibitory action of infrared stimulation.

Our pharmacological results support this interpretation. Blocking GABA_A_ receptors with picrotoxin under anesthesia shifted INS responses toward the negative pattern observed during wakefulness, whereas enhancing GABA_A_ receptors with muscimol in awake animals shifted responses toward the positive pattern observed under anesthesia. At first sight, these effects may appear paradoxical if INS is assumed to act simply by increasing or decreasing inhibition. Instead, they suggest that the effect of INS depends on the pre-existing excitatory-inhibitory configuration of the brain circuit. Both drugs alter not only GABAergic transmission but also spontaneous activity and network excitability; therefore, these experiments do not establish a specific inhibitory pathway as the mechanism of INS. Rather, together with the baseline-dependence analysis, they support a model in which the same external input is transformed differently according to the physiological state of the cortical network. A feed-forward inhibitory circuit provides one plausible framework for this state-dependent transformation, but direct measurements of membrane potential, synaptic excitation and inhibition, and cell-type-specific activity will be required to test this mechanism.

### Integration of INS with ongoing neural activity

The broader implication of these findings is that focal neural stimulation should not necessarily be viewed as an external command that simply overrides ongoing activity. Instead, its neuronal effect emerges from an interaction between the imposed stimulus and the dynamical brain state of the circuit receiving it^48,49^. In this framework, stimulation intensity and spatial targeting define the external input, whereas membrane potential, spontaneous activity, excitation-inhibition balance, and network connectivity define the physiological context in which that input is transformed. The same physical stimulus can therefore produce substantially different, and in our experiments, opposite neuronal responses without any change in stimulation parameters.

This view is particularly relevant to INS because its high spatial focality has generally been considered one of its principal advantages for precise neuromodulation. Spatial precision, however, does not necessarily confer functional predictability. Our results suggest that even when the same restricted neuronal population is targeted with the same stimulation parameters, the resulting response can change with physiological state. Consequently, optimization of INS may require consideration not only of radiant exposure, pulse pattern, and fiber geometry, but also of the ongoing brain state of the targeted circuit.

Such state dependence also provides a rationale for future closed-loop implementations of INS. Rather than delivering stimulation according to fixed parameters, adaptive systems could incorporate measurements of ongoing neural activity to select stimulation conditions appropriate for the current circuit state. Similar principles underlie adaptive deep brain stimulation, brain-computer interfaces, and other feedback-guided neuromodulation approaches^56–60^. INS may be particularly suited to such strategies because its focality permits spatially restricted stimulation while physiological signals provide complementary information about when and how that stimulation should be delivered. Establishing reliable physiological biomarkers of INS responsiveness and determining whether the state dependence observed here generalizes across cortical areas, behavioral conditions, and species will be important next steps toward context-aware neuromodulation.

## Acknowledgments

We thank Yousheng Shu and Lang Wang for discussions and comments on the manuscript. This work was supported by the grants from National Key R&D program of China Brain Initiative 2021ZD0200401 (A.W.R.; W.X.); National Natural Science Foundation of China U20A20221 (A.W.R.), 81961128029 (A.W.R.), 31627802 (A.W.R.); Key Research and Development Program of Zhejiang Province 2022C03096 (W.X.), 2024SSYS0019 (A.W.R.); Fundamental Research Funds for the Central Universities 226-2022-00083 (W.X.); National Natural Science Foundation of China 82602376 (P.F.); Health Science and Technology Project of Hubei Province WJ2025Q099 (P.F.); Scientific Research Plan of the Education Department of Hubei Province Q20251111 (P.F.).

## Author contributions

Conceptualization, A.W.R., W.X., P.F.; Methodology, P.F., Y.L., H.Z., W.X., A.W.R.; Formal Analysis, P.F., W.Z., W.X., A.W.R.; Investigation, P.F., Y.L., W.X., A.W.R.; Writing – Original Draft, P.F., W.X., A.W.R.; Writing – Review & Editing, P.F., W.X., A.W.R., Y.Z., H.Z., L.Z., M.W., Y.Y., S.S.; Visualization, P.F., W.X., A.W.R., S.S., L.Z., Y.P.; Supervision, W.X., A.W.R., Y.Z.; Project Administration, W.X., A.W.R., Y.Z., P.F., Y.L.; Funding Acquisition, W.X., A.W.R., P.F.

## Methods

### Surgery

All procedures were approved by the Zhejiang University Animal Experimentation Committee and were completed in compliance with the National Institutes of Health Guide for the Care and Use of Laboratory Animals. The surgical procedures used in these experiments have been described previously^14^ and are briefly described below.

Female C57BL/6J mice (8-10 weeks old) were group-housed (3 per cage) on a 12 h light-dark cycle and provided with food and water *ad libitum*. Each mouse underwent a total of three separate surgical procedures to inject 1) AAV virus, and chronically implant 2) a glass cranial window and 3) a headplate. Mice were anesthetized with isoflurane (5% inhalation, mixed with fresh air, 0.5 L/min) and placed in a stereotactic frame, after which isoflurane was maintained at 2% throughout the surgical procedures, with body temperature maintained at 37 ℃ by a heating pad.

A craniotomy was performed by measuring 4 mm in diameter, exposing the somatosensory cortex (at coordinates of 0.3 mm posterior and 2.3 mm lateral from bregma). The dura was carefully removed, and mice were injected with 200 nL of virus (neuron: rAVV2/9-hSyn-GCaMP6s-WPRE-hGH-pA, titer = 1.03×10^13^ vg/ml, PT-0145, BrainVTA; rAVV2/9-hSyn-EGFP-WPRE-hGH-pA, titer = 5.5×10^12^ vg/ml, PT-1990, BrainVTA) into 3-5 locations at 400 μm below the cortex pia by a patch pipette. A coverglass (measuring 6 mm in diameter) was used to cover the brain cortex and secured with medical glue at coverglass edge. A custom-made headplate was glued to the skull with dental acrylic to fill the gap between the headplate and the skull, allowing the mice to be head-fixed while imaged awake. A chronic imaging window was completed, which can be used repeatedly to immobilize in the same position in both awake and anesthetized mice. Mice received an intraperitoneal injection of ceftriaxone (2.5 mg/kg) and a subcutaneous injection of buprenorphine (0.05 mg/kg) for up to three days post-surgery and were allowed to recover for at least three weeks before they were imaged.

### Infrared neural modulation

A customized single-wavelength laser (FC-W series, CNI) coupled with multimode fiber (MM200, Newdoon) was used to deliver a 1,875 nm infrared beam. The fiber (200 μm core diameter, 0.37 NA) was positioned with a custom-built micromanipulator at an angle of approximately 45° to the brain surface (Fig. 1b). The pulse energy was calibrated with a power meter (FieldMaxII-TO, Coherent Inc.) before each experiment and transmitted through the laser fiber. Seven radiant exposures (0.16, 0.29, 0.42, 0.50, 0.59, 0.68, 0.76 J/cm^2^ per pulse, see reference^14^) were presented in pseudo-random order to prevent trends, with six trials per intensity. Each 60-s trial contained a 10-s pre-stimulation baseline, six 0.5-s INS pulse trains separated by 2.5 s, and a 32-s post-stimulation recovery period. Each train consisted of 100 pulses delivered at 200 Hz with a pulse width of 0.25 ms (Fig. 1d). The two-photon system generated a TTL trigger to synchronize INS delivery with imaging.

As described in our previous study^14^, we modeled tissues in a cube of 4 mm slide length to simulate light propagation in planar multi-layered tissues. We modified the codes for our infrared light (λ = 1,875 nm) propagation in multi-layered tissues^61,62^ and then defined the illuminated region for focal effect.

### Two-photon calcium imaging

Calcium responses of cortical neurons were acquired using a customized two-photon microscope (Ultima IV, Bruker Corporation) coupled with a femtosecond mode-locked Ti: Sapphire laser (80 MHz, 140 fs, Chameleon Ultra II, Coherent Inc.). The femtosecond laser was set for GCaMP6s or EGFP excitation at a wavelength of 920 nm. A Pockels Cell (EO-PC, Thorlabs Corporation) was used to regulate laser power. For calcium imaging, laser power after the 16X objective (N16XLWD-PF, 0.8 NA, Nikon) was limited to a maximum of 40 mW, depending on depth. Emission light was filtered using a bandpass 525/70 filter for GCaMP6s or EGFP and measured by GaAsP photomultiplier tubes (model H10770, Hamamatsu Photonics). Imaging sequences were collected at ∼30 Hz using a resonant-galvo scanner with 512×512 pixel resolution for functional imaging. Images were collected using a galvo-galvo scanner with 1024×1024 pixel resolution for structural imaging. For awake experiments with INS application, three days of routine handling of the mice were used to acclimate them to the imaging system, and the immobilization device greatly helped to reduce animal motion. For anesthetized experiments with INS application, isoflurane (0.6%, mixed with fresh air, 0.5 L/min) was used to anesthetize mice, with body temperature maintained by a heating pad during imaging.

### Pharmacology

For pharmacological experiments, the original coverglass was replaced with a 5-mm-diameter coverglass containing a 500-μm central opening to permit local drug application. Picrotoxin (PTX, product number B5054, APExBIO) was applied in anesthetized mice to mimick an awake-like state, while muscimol (MSM, product number 5.06044, Sigma-Aldrich) was applied in awake mice to mimick an anesthetized-like state (Supplementary Fig. 5). Spontaneous neuronal calcium activity in the anesthetized or awake state was first obtained, and then pharmacological experiments were performed. For PTX experiments, artificial cerebrospinal fluid (ACSF) was replaced with PTX at a concentration of 100 μM, and incubated for two hours. For MSM experiments, ACSF was replaced with MSM at a concentration of 100 μM, and incubated for two hours. After incubation, the solution was replaced with ACSF, after which spontaneous activity and INS-evoked calcium activity were recorded. These manipulations were used to alter GABA_A_ receptor signaling and network state; they were not interpreted as selective manipulations of baseline calcium activity alone.

### Data analysis

#### Registration, motion correction, cell detection, and calcium extraction

Acquired time-lapse two-photon images were read and analyzed in FIJI (Fiji Is Just ImageJ, NIH), Cellpose (version 2.0), and MATLAB (version R2020b, MathWorks). Image registration in the xy plane was corrected by using the template_matching plugin in FIJI^63,64^. Cellpose was used to detect neuronal soma except neuropil, and perform cellular segmentation to get masks of the region of interest (ROI)^65^. FIJI was then used to register raw movies, use masks, and extract calcium fluorescence traces. Raw fluorescence (F) was calculated by averaging the corresponding pixel values in each specified ROI mask. In the following analysis, we used our self-written MATLAB (version R2020b, MathWorks) code. Relative fluorescence changes were calculated as ΔF/F_0_ = (F-F_0_)/F_0_ for each trial and neuron, with F_0_ defined as the mean fluorescence during the first 10 s before stimulation onset (pre-INS). The calculated average of six trials for each neuron was used as the mean activity for that neuron (see Supplementary Fig. 4), as shown in each row in the heatmaps at different laser intensities. Only xy motion correction was applied; thus, possible residual out-of-plane motion is an acknowledged limitation of awake imaging and is considered together with the temporal, spatial, pharmacological, and EGFP controls when interpreting negative fluorescence deflections.

#### Quantification of spontaneous calcium activity

To quantify the spontaneous activity, we first combined the six spontaneous trials into a single timecourse and identified the calcium peaks. We then employed the MATLAB ‘findpeaks’ function with the following criterion: ΔF/F_0_ exceeding 0.1 and lasting for at least 1 s. Finally, we calculated two indices: 1) peak counts, the number of calcium peaks; 2) peak duration, the total lasting time of all calcium peaks. In awake, anesthetized, and pharmacological experiments, we measured spontaneous calcium activity using these two indices.

#### Descriptive stratification and quantification of INS-evoked neuronal responses

INS-evoked responses were quantified from somatic fluorescence relative to the pre-stimulus period. For each neuron and intensity, response amplitude was calculated from the mean ΔF/F_0_ across the six stimulation-train periods (pt1, pt2, pt3, pt4, pt5, pt6; mean ΔF/F_0_, Supplementary Fig. 4). To describe heterogeneity in awake-to-anesthetized response changes, the averaged responses of the paired neurons were stratified using MATLAB k-means clustering at each intensity (‘kmeans’ function; the number of clusters k = 4, replicates of the kmeans process = 10). The resulting clusters were subsequently grouped according to the direction of the response change (increasing, relatively unchanged, or decreasing). This procedure was used as an exploratory, analysis-defined stratification of response patterns and was not intended to identify biological neuronal subclasses. Response-defined groups were then used for descriptive intensity-dependence and pre-stimulus association analyses.

#### In situ calibration of INS–induced thermal artifacts

To estimate the direct fluoro-thermal contribution of pulsed INS to fluorescence measurements, we performed a calibration experiment using cortical neurons expressing EGFP. Because GCaMP indicators are derived from GFP-family fluorophores, EGFP provides a calcium-insensitive reference for temperature-related fluorescence changes under matched optical stimulation conditions. Adult mice (*n* = 3) expressing AAV–hSyn–EGFP in the somatosensory cortex were imaged while awake and head-fixed during INS at 0.16, 0.42, 0.59, and 0.76 J/cm^2^. For each neuron, the fluorescence timecourse was normalized to the pre–stimulus baseline, and the calcium deflection (ΔF/F_0_) was extracted per trial. To characterize the spatial profile of heating, single-cell coordinates from the three animals were normalized to the fiber-tip position (set as the origin). Two–dimensional spatial maps (Fig. 7f and 7g) were generated by applying Gaussian process regression model with a Kriging kernel to the single–cell -ΔF/F_0_ values; the mean -ΔF/F_0_ was then binned by radial distance from the tip (0-25, 25-50, 50-75, 75-100, and >100 μm) for each intensity.

For cross–calibration with GCaMP6s recordings (performed in separate cohorts under identical INS parameters), we compared the amplitudes (ΔF/F_0_) of raw GCaMP6s responses with the corresponding calibration results of subtracting EGFP thermal measurements to account for differences in temperature sensitivity. Certainly, in this comparison, the EGFP measurements were used as an empirical population-level estimate of the expected thermal fluorescence component. Because the reporters were measured in different animals, this analysis was used to estimate, rather than directly subtract on a cell-by-cell basis, the contribution that could plausibly be attributed to fluorophore heating.

To further quantify the intensity-dependence of the fluoro-thermal contribution, we calculated an EGFP-based thermal shift for each radiant exposure. The thermal shift was defined as the absolute difference between the calibrated and uncorrected GCaMP6s response amplitudes: Thermal shift = |R_calibration_ - R_raw_|, where R_raw_ represents the measured GCaMP6s response amplitude and R_calibration_ represents the corresponding response after applying the EGFP-derived thermal measurements. Thermal shifts were calculated for 0.16, 0.42, 0.59, and 0.76 J/cm^2^. To compare lower and higher radiant exposures, the mean thermal shift at 0.59 and 0.76 J/cm^2^ was divided by the mean thermal shift at 0.16 and 0.42 J/cm^2^, yielding the 3.37-fold increase in thermal influence between the two exposure ranges. In addition, a relative thermal influence (RTI) index was calculated by normalizing the thermal shift at each radiant exposure to the maximal thermal shift observed at 0.76 J/cm^2^: RTI (%) = Thermal shift_intensity_ / Thermal shift_0.76J/cm2_ × 100%. This normalization provides an intensity-dependent measure of the relative magnitude of the EGFP-estimated fluoro-thermal component, with the response at 0.76 J/cm^2^ defined as 100%. Because the EGFP calibration was derived independently of physiological state and applied equivalently to awake and anesthetized datasets, these metrics quantify the intensity dependence of the estimated thermal fluorescence contribution rather than a state-specific thermal response.

### Statistical analysis

Data are presented as mean ± SEM unless otherwise specified; timecourse traces are shown as mean ± SD. Paired Wilcoxon tests were used for within-neuron or within-pair comparisons in Fig. 2h, Fig. 3b, Fig. 4c, Fig. 5c, Fig. 6a, Fig. 6e, Supplementary Fig. 2b, 2c, Supplementary Fig. 3b, and Supplementary Fig. 7. Mann-Whitney tests were used for unpaired comparisons in Fig. 7i, Supplementary Fig. 2a, and Supplementary Fig. 6b, 6d. Linear regressions were used for the dose-level and trial-by-trial association analyses reported in the corresponding figures and supplementary tables. The exact *n*, comparison, and *p* value for each analysis are specified in the figure legends or tables. In all tests, *p* < 0.05 is considered statistically significant, and ‘ns’ represents no significance. Statistical analyses were performed in Prism (version 8.0, GraphPad Software). The analyses were conducted at the neuron or trial level as indicated, because repeated measurements are nested within a small number of animals. No additional methods were used to determine whether the data met the assumptions of the statistical approach. The statistical details of each experiment can be found in the figure legends and tables.

## Reference

1 Bradley, C., Nydam, A. S., Dux, P. E. & Mattingley, J. B. State-dependent effects of neural stimulation on brain function and cognition. Nat Rev Neurosci 23, 459–475, doi:10.1038/s41583-022-00598-1 (2022).

2 Guidetti, M. et al. Clinical perspectives of adaptive deep brain stimulation. Brain Stimul 14, 1238–1247, doi:10.1016/j.brs.2021.07.063 (2021).

3 Griffin, D. M., Hudson, H. M., Belhaj-Saïf, A. & Cheney, P. D. Hijacking cortical motor output with repetitive microstimulation. J Neurosci 31, 13088–13096, doi:10.1523/jneurosci.6322-10.2011 (2011).

4 Cheney, P. D., Griffin, D. M. & Van Acker, G. M., 3rd. Neural hijacking: action of high-frequency electrical stimulation on cortical circuits. Neuroscientist 19, 434–441, doi:10.1177/1073858412458368 (2013).

5 Neumann, W. J., Steiner, L. A. & Milosevic, L. Neurophysiological mechanisms of deep brain stimulation across spatiotemporal resolutions. Brain 146, 4456–4468, doi:10.1093/brain/awad239 (2023).

6 Balbinot, G. et al. The mechanisms of electrical neuromodulation. J Physiol 603, 247–284, doi:10.1113/jp286205 (2025).

7 Chernov, M. & Roe, A. W. Infrared neural stimulation: a new stimulation tool for central nervous system applications. Neurophotonics 1, 011011, doi:10.1117/1.NPh.1.1.011011 (2014).

8 Ping, A. et al. Targeted optical neural stimulation: a new era for personalized medicine. Neuroscientist, doi:10.1177/10738584211057047 (2021).

9 Jiang, S., Wu, X., Rommelfanger, N. J., Ou, Z. & Hong, G. Shedding light on neurons: optical approaches for neuromodulation. Natl Sci Rev 9, nwac007, doi:10.1093/nsr/nwac007 (2022).

10 LaLumiere, R. T. A new technique for controlling the brain: optogenetics and its potential for use in research and the clinic. Brain Stimul 4, 1–6, doi:10.1016/j.brs.2010.09.009 (2011).

11 Lüscher, C. et al. Roadmap for direct and indirect translation of optogenetics into discoveries and therapies for humans. Nat Neurosci 28, 2415–2431, doi:10.1038/s41593-025-02097-9 (2025).

12 Chernov, M. M., Chen, G. & Roe, A. W. Histological assessment of thermal damage in the brain following infrared neural stimulation. Brain Stimul 7, 476–482, doi:10.1016/j.brs.2014.01.006 (2014).

13 Pan, L. et al. Infrared neural stimulation in human cerebral cortex. Brain Stimul 16, 418–430, doi:10.1016/j.brs.2023.01.1678 (2023).

14 Fu, P. et al. Two-photon imaging of excitatory and inhibitory neural response to infrared neural stimulation. Neurophotonics 11, 025003, doi:10.1117/1.NPh.11.2.025003 (2024).

15 Cayce, J. M., Friedman, R. M., Jansen, E. D., Mahavaden-Jansen, A. & Roe, A. W. Pulsed infrared light alters neural activity in rat somatosensory cortex in vivo. Neuroimage 57, 155–166, doi:10.1016/j.neuroimage.2011.03.084 (2011).

16 Cayce, J. M. et al. Calcium imaging of infrared-stimulated activity in rodent brain. Cell Calcium 55, 183–190, doi:10.1016/j.ceca.2014.01.004 (2014).

17 Tian, F., Zhang, Y., Schriver, K. E., Hu, J. M. & Roe, A. W. A novel interface for cortical columnar neuromodulation with multipoint infrared neural stimulation. Nat Commun 15, 6528, doi:10.1038/s41467-024-50375-0 (2024).

18 Cayce, J. M. et al. Infrared neural stimulation of primary visual cortex in non-human primates. Neuroimage 84, 181–190, doi:10.1016/j.neuroimage.2013.08.040 (2014).

19 Xu, A. G. et al. Focal infrared neural stimulation with high-field functional MRI: a rapid way to map mesoscale brain connectomes. Sci Adv 5, eaau7046, doi:10.1126/sciadv.aau7046 (2019).

20 Chernov, M. M., Friedman, R. M. & Roe, A. W. Fiberoptic array for multiple channel infrared neural stimulation of the brain. Neurophotonics 8, 025005, doi:10.1117/1.NPh.8.2.025005 (2021).

21 Tian, F., Liu, Y., Chen, M., Schriver, K. E. & Roe, A. W. Selective activation of mesoscale functional circuits via multichannel infrared stimulation of cortical columns in ultra-high-field 7T MRI. Cell Rep Methods 5, 100961, doi:10.1016/j.crmeth.2024.100961 (2025).

22 Ping, A. et al. Brainwide mesoscale functional networks revealed by focal infrared neural stimulation of the amygdala. Natl Sci Rev 12, nwae473, doi:10.1093/nsr/nwae473 (2025).

23 Shi, S. H. et al. Infrared neural stimulation with 7T fMRI: a rapid in vivo method for mapping cortical connections of primate amygdala. Neuroimage 231, 117818, doi:10.1016/j.neuroimage.2021.117818 (2021).

24 Yao, S. et al. Functional topography of pulvinar-visual cortex networks in macaques revealed by INS-fMRI. J Comp Neurol 531, 681–700, doi:10.1002/cne.25456 (2023).

25 Feng, Y. et al. Infrared neural stimulation with fMRI in primates reveals mesoscale limbic organization linked to the medial pulvinar. Cell Rep 45, 117209, doi:10.1016/j.celrep.2026.117209 (2026).

26 Kaszas, A. et al. Two-photon GCaMP6f imaging of infrared neural stimulation evoked calcium signals in mouse cortical neurons in vivo. Sci Rep 11, 9775, doi:10.1038/s41598-021-89163-x (2021).

27 Dadarlat, M. C., Sun, Y. J. & Stryker, M. P. Activity-dependent recruitment of inhibition and excitation in the awake mammalian cortex during electrical stimulation. Neuron 112, 821–834.e824, doi:10.1016/j.neuron.2023.11.022 (2024).

28 Pasley, B. N., Allen, E. A. & Freeman, R. D. State-dependent variability of neuronal responses to transcranial magnetic stimulation of the visual cortex. Neuron 62, 291–303, doi:10.1016/j.neuron.2009.03.012 (2009).

29 Lothet, E. H. et al. Selective inhibition of small-diameter axons using infrared light. Sci Rep 7, 3275, doi:10.1038/s41598-017-03374-9 (2017).

30 Fu, P., et al. Two-photon imaging of GABAergic and non-GABAergic neuronal calcium activity induced by infrared neural stimulation in awake mouse cortex. SPIE BiOS 12366 (2023).

31 Plaksin, M., Shapira, E., Kimmel, E. & Shoham, S. Thermal transients excite neurons through universal intramembrane mechanoelectrical effects. Phys Rev X 8, doi:10.1103/PhysRevX.8.011043 (2018).

32 Shapiro, M. G., Homma, K., Villarreal, S., Richter, C. P. & Bezanilla, F. Infrared light excites cells by changing their electrical capacitance. Nat Commun 3, 736, doi:10.1038/ncomms1742 (2012).

33 Ali, F. & Kwan, A. C. Interpreting in vivo calcium signals from neuronal cell bodies, axons, and dendrites: a review. Neurophotonics 7, 011402, doi:10.1117/1.NPh.7.1.011402 (2020).

34 Forli, A. et al. Two-photon bidirectional control and imaging of neuronal excitability with high spatial resolution in vivo. Cell Rep 22, 3087–3098, doi:10.1016/j.celrep.2018.02.063 (2018).

35 Vogt, K. E., Gerharz, S., Graham, J. & Canepari, M. High-resolution simultaneous voltage and Ca^2+^ imaging. J Physiol-London 589, 489–494, doi:10.1113/jphysiol.2010.200220 (2011).

36 Voss, L. J. & Garcia, V. Electrophysiological field potential identification of an intact GABAergic system in mouse cortical slices. Brain Res 1756, doi:10.1016/j.brainres.2021.147295 (2021).

37 Liu, Y. U. et al. Neuronal network activity controls microglial process surveillance in awake mice via norepinephrine signaling. Nature Neuroscience 22, 1771–1781, doi:10.1038/s41593-019-0511-3 (2019).

38 Akk, G. et al. Enhancement of muscimol binding and gating by allosteric modulators of the GABA_A_ receptor: relating occupancy to state functions. Mol Pharmacol 98, 303–313, doi:10.1124/molpharm.120.000066 (2020).

39 Zhang, X. Q., Yao, N. & Chergui, K. The GABA(A) receptor agonist muscimol induces an age- and region-dependent form of long-term depression in the mouse striatum. Learn Memory 23, 479–485, doi:10.1101/lm.043190.116 (2016).

40 Estrada, H. et al. High-resolution fluorescence-guided transcranial ultrasound mapping in the live mouse brain. Sci Adv 7, doi:10.1126/sciadv.abi5464 (2021).

41 Kamei, Y. et al. Infrared laser-mediated gene induction in targeted single cells in vivo. Nat Methods 6, 79–81, doi:10.1038/nmeth.1278 (2009).

42 Roe, A. W. et al. Infrared neural stimulation in V1 produces intensity-dependent perceptual biases in awake macaques. SPIE Vol. 13836 (2026).

43 Xi, Y., Schriver, K. E., Roe, A. W. & Zhang, X. Quantifying tissue temperature changes induced by infrared neural stimulation: numerical simulation and MR thermometry. Biomed Opt Express 15, 4111–4131, doi:10.1364/boe.530854 (2024).

44 Treviño, M. Inhibition controls asynchronous states of neuronal networks. Front Synaptic Neurosci 8, 11, doi:10.3389/fnsyn.2016.00011 (2016).

45 Ishizuka, N., Cowan, W. M. & Amaral, D. G. A quantitative analysis of the dendritic organization of pyramidal cells in the rat hippocampus. J Comp Neurol 362, 17–45, doi:10.1002/cne.903620103 (1995).

46 Amaral, D. G. & Witter, M. P. The three-dimensional organization of the hippocampal formation: a review of anatomical data. Neuroscience 31, 571–591, doi:10.1016/0306-4522(89)90424-7 (1989).

47 Higley, M. J. & Contreras, D. Balanced excitation and inhibition determine spike timing during frequency adaptation. J Neurosci 26, 448–457, doi:10.1523/Jneurosci.3506-05.2006 (2006).

48 Murris, S. R., Arsenault, J. T. & Vanduffel, W. Frequency- and State-Dependent Network Effects of Electrical Stimulation Targeting the Ventral Tegmental Area in Macaques. Cereb Cortex 30, 4281–4296, doi:10.1093/cercor/bhaa007 (2020).

49 Yang, Y. et al. Modelling and prediction of the dynamic responses of large-scale brain networks during direct electrical stimulation. Nat Biomed Eng 5, 324–345, doi:10.1038/s41551-020-00666-w (2021).

50 Kozyrev, V., Staadt, R., Eysel, U. T. & Jancke, D. TMS-induced neuronal plasticity enables targeted remodeling of visual cortical maps. Proc Natl Acad Sci U S A 115, 6476–6481, doi:10.1073/pnas.1802798115 (2018).

51 Grosshagauer, S. et al. Chronometric TMS-fMRI of personalized left dorsolateral prefrontal target reveals state-dependency of subgenual anterior cingulate cortex effects. Mol Psychiatry 29, 2678–2688, doi:10.1038/s41380-024-02535-3 (2024).

52 Janssens, S. E. W. & Sack, A. T. Spontaneous fluctuations in oscillatory brain state cause differences in transcranial magnetic stimulation effects within and between individuals. Front Hum Neurosci 15, 802244, doi:10.3389/fnhum.2021.802244 (2021).

53 Oliver, A. E., Baker, G. A., Fugate, R. D., Tablin, F. & Crowe, J. H. Effects of temperature on calcium-sensitive fluorescent probes. Biophys J 78, 2116–2126, doi:10.1016/S0006-3495(00)76758-0 (2000).

54 Feng, H. J. et al. Alteration of GABAergic neurotransmission by pulsed infrared laser stimulation. J Neurosci Meth 192, 110–114, doi:10.1016/j.jneumeth.2010.07.014 (2010).

55 Zhu, X., Lin, J. W., Turnali, A. & Sander, M. Y. Single infrared light pulses induce excitatory and inhibitory neuromodulation. Biomed Opt Express 13, 374–388, doi:10.1364/BOE.444577 (2022).

56 Wang, X., et al. A longitudinal electrophysiological and behavior dataset for PD rat in response to deep brain stimulation. Sci Data 11, 500, doi:10.1038/s41597-024-03356-3 (2024).

57 Amoozegar, S., Pooyan, M. & Roughani, M. Toward a closed-loop deep brain stimulation in Parkinson’s disease using local field potential in parkinsonian rat model. Med Hypotheses 132, 109360, doi:10.1016/j.mehy.2019.109360 (2019).

58 Coventry, B. S., Lawlor, G. L., Bagnati, C. B., Krogmeier, C. & Bartlett, E. L. Characterization and closed-loop control of infrared thalamocortical stimulation produces spatially constrained single-unit responses. PNAS Nexus 3, pgae082, doi:10.1093/pnasnexus/pgae082 (2024).

59 Shanechi, M. M. Brain-machine interfaces from motor to mood. Nat Neurosci 22, 1554–1564, doi:10.1038/s41593-019-0488-y (2019).

60 Sun, G. et al. Closed-loop stimulation using a multiregion brain-machine interface has analgesic effects in rodents. Sci Transl Med 14, eabm5868, doi:10.1126/scitranslmed.abm5868 (2022).

61 Wang, L., Jacques, S. L. & Zheng, L. MCML--Monte Carlo modeling of light transport in multi-layered tissues. Comput Methods Programs Biomed 47, 131–146, doi:10.1016/0169-2607(95)01640-f (1995).

62 Shin, Y. & Kwon, H. S. Mesh-based Monte Carlo method for fibre-optic optogenetic neural stimulation with direct photon flux recording strategy. Phys Med Biol 61, 2265–2282, doi:10.1088/0031-9155/61/6/2265 (2016).

63 Sharma, K., Gordon, G. R. J. & Tran, C. H. T. Heterogeneity of sensory-induced astrocytic Ca^2+^ dynamics during functional hyperemia. Front Physiol 11, 611884, doi:10.3389/fphys.2020.611884 (2020).

64 Tran, C. H. T., Peringod, G. & Gordon, G. R. Astrocytes integrate behavioral state and vascular signals during functional hyperemia. Neuron 100, 1133–1148, doi:10.1016/j.neuron.2018.09.045 (2018).

65 Stringer, C., Wang, T., Michaelos, M. & Pachitariu, M. Cellpose: a generalist algorithm for cellular segmentation. Nat Methods 18, 100–106, doi:10.1038/s41592-020-01018-x (2021).

