## Supplemental information for "Brain state shapes the polarity of pulsed infrared neural stimulation responses in individual cortical neurons"

### Supplementary Figures and Tables

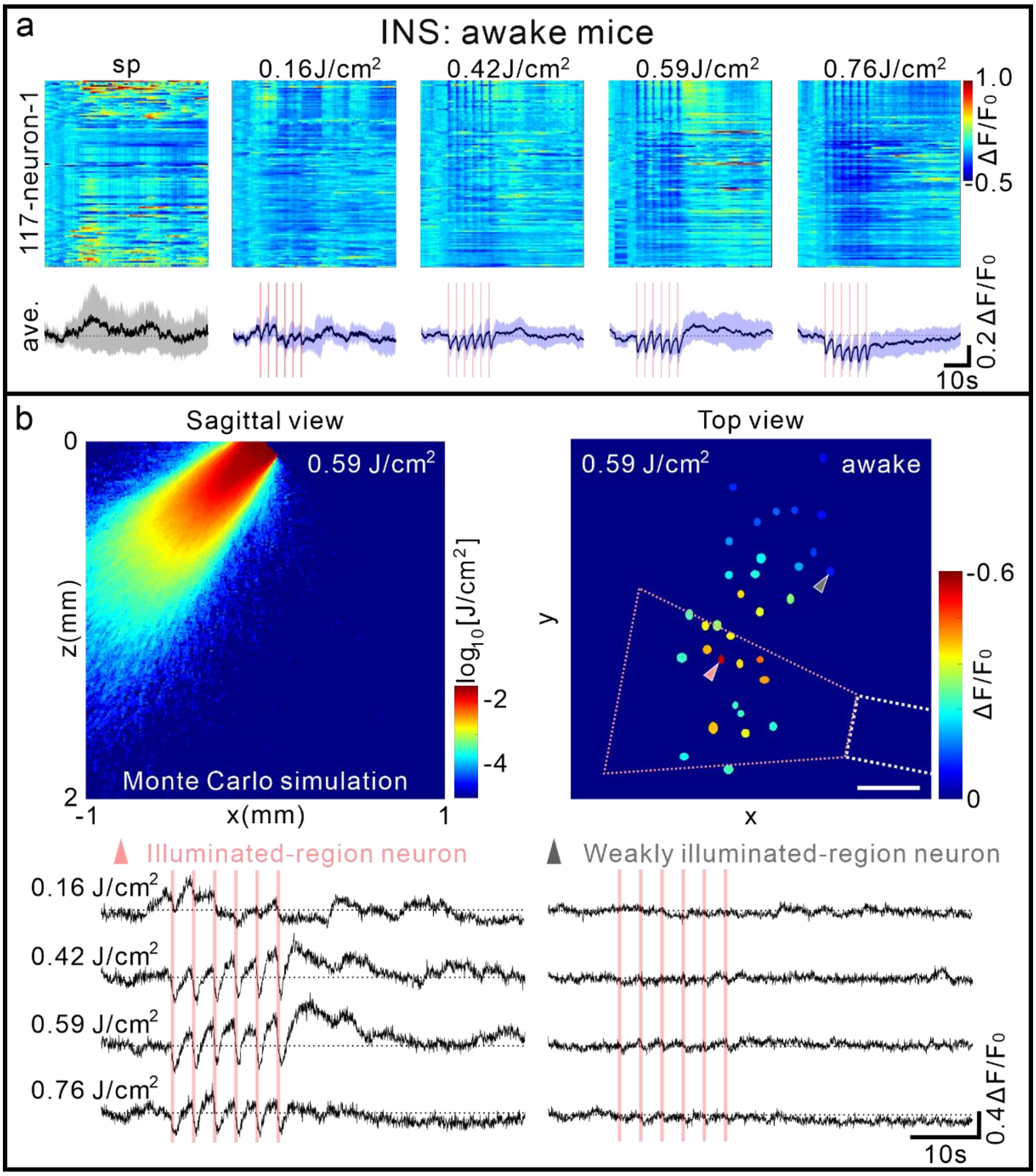


**Supplementary Fig. 1 INS elicits negative responses and exerts focal modulatory effects in awake mice.**

(a) INS induces consistent negative neuronal responses in awake mice in an independent dataset. The pink bar denotes the INS application period, and shaded areas represent the standard deviation (SD). (b) Focal calcium response profiles upon INS stimulation in awake mice. Neurons within the light-illuminated region exhibit stronger INS-evoked responses across all four stimulation intensities, which is consistent with our previous observations in anesthetized mice. Related to Figs. 1 and 2.

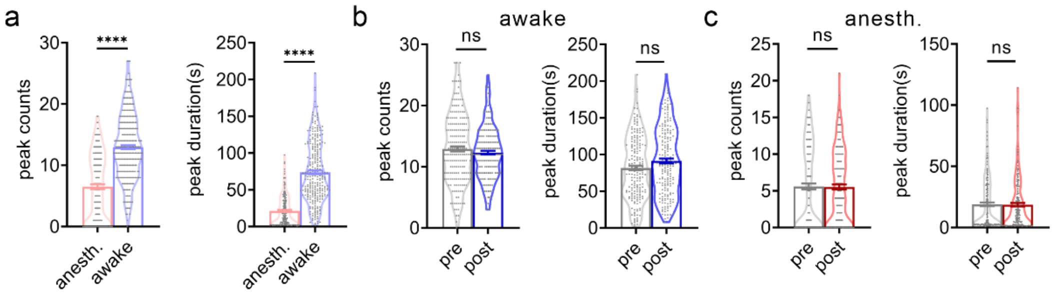


**Supplementary Fig. 2 Spontaneous neuronal calcium activity in awake and anesthetized mice.** (a) Differences in spontaneous calcium activity between anesthetized and awake mice. Peak count: 6.50 ± 0.38 (anesthetized) versus 12.96 ± 0.32 (awake); peak duration: 21.18 ± 1.43 s (anesthetized) versus 73.80 ± 2.48 s (awake). Unpaired Mann–Whitney test, $p < 0.0001$ for both metrics. (b) Changes in spontaneous calcium activity before and after INS stimulation in awake mice. Peak count: 12.92 ± 0.42 (pre-INS) vs. 12.30 ± 0.32 (post-INS), paired Wilcoxon test, $p = 0.3252$; peak duration: 81.58 ± 3.14 s (pre-INS) vs. 91.27 ± 3.37 s (post-INS), paired Wilcoxon test, $p = 0.0597$. (c) Spontaneous calcium activity before and after INS stimulation in anesthetized mice (modified from Ref. 14). Peak count: 5.59 ± 0.41 (pre-INS) vs. 5.51 ± 0.35 (post-INS), paired Wilcoxon test, $p = 0.5180$; peak duration: 18.97 ± 1.49 s (pre-INS) vs. 18.61 ± 1.46 s (post-INS), paired Wilcoxon test, $p = 0.6488$. Data are presented as mean ± SEM. Statistical analyses were performed using the Wilcoxon test. anesth., anesthetized. Related to Fig. 2.

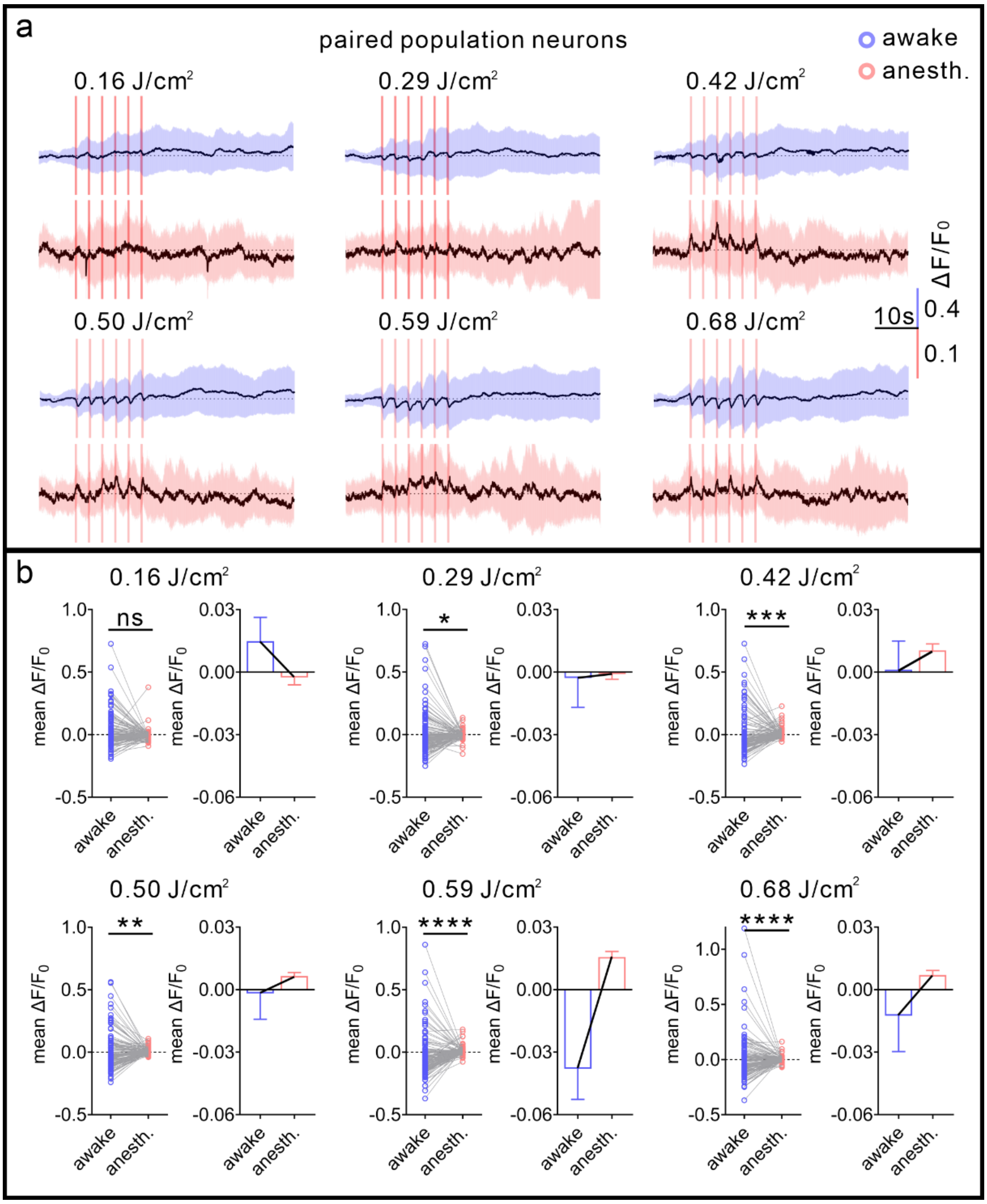


**Supplementary Fig. 3 Paired neuronal population responses under awake and anesthetized conditions.**

(a) INS elicits negative responses in neurons from awake mice, whereas their paired counterparts exhibit excitatory responses under anesthesia. The pink bar marks the INS application period; shaded areas denote the standard deviation (SD). (b) Mean response amplitude during INS stimulation across six stimulation intensities in the awake and anesthetized states (paired Wilcoxon test). Data are presented as mean ± SEM. Related to Fig. 2.

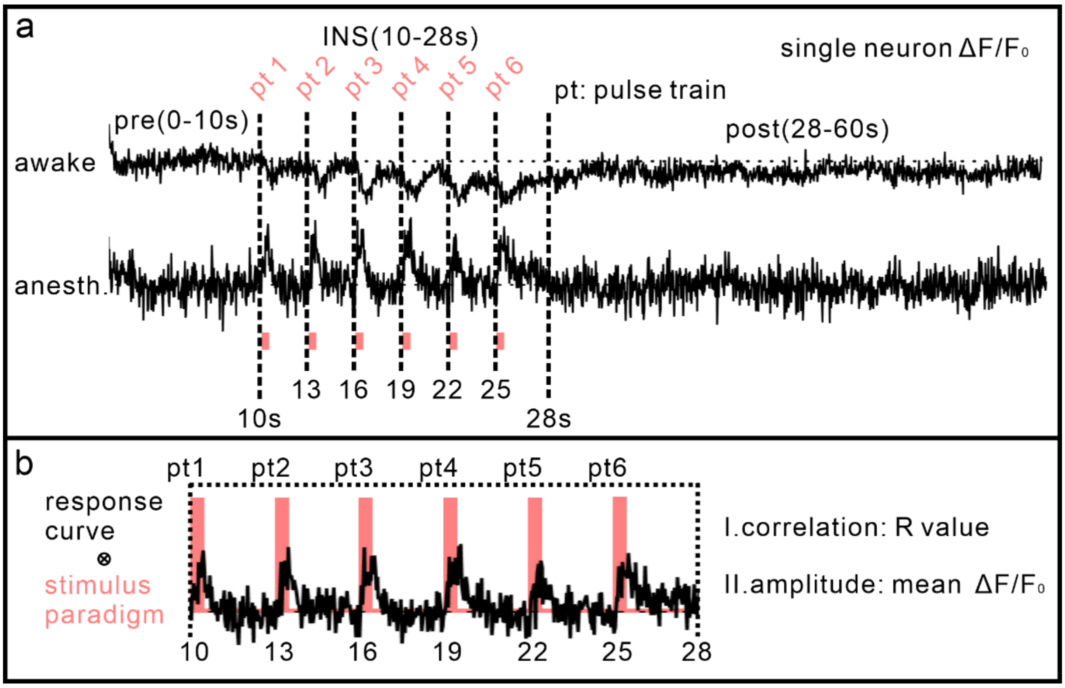


**Supplementary Fig. 4 Schematic illustration of the quantitative index calculation for INS-evoked neuronal responses.**

(a) A 60-s recording time course divided into three epochs: baseline (pre-INS, 0–10 s), INS stimulation (10–28 s), and post-stimulation (post-INS, 28–60 s), for the individual neuron recorded under both awake and anesthetized conditions. (b) Calculation of response correlation and amplitude for each neuron across six repeated pulse train periods (pt1–pt6) within the INS stimulation window. Related to Fig. 2.

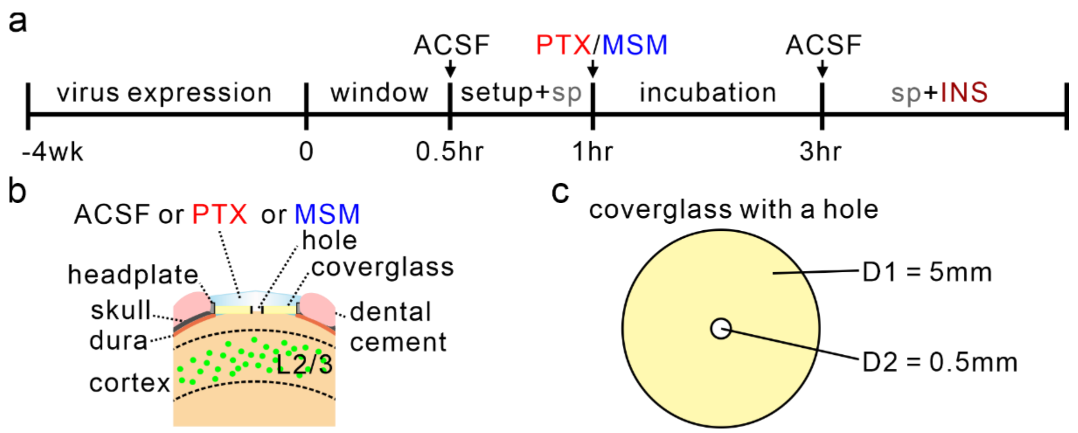


**Supplementary Fig. 5 Schematic illustration of drug incubation procedures and perforated coverglass setup.**

(a) Experimental timeline for drug incubation and related manipulations. (b) Schematic of the cranial window setup for two-photon imaging. (c) Dimensional schematic of the custom-manufactured perforated coverglass. Related to Figs. 4 and 5.

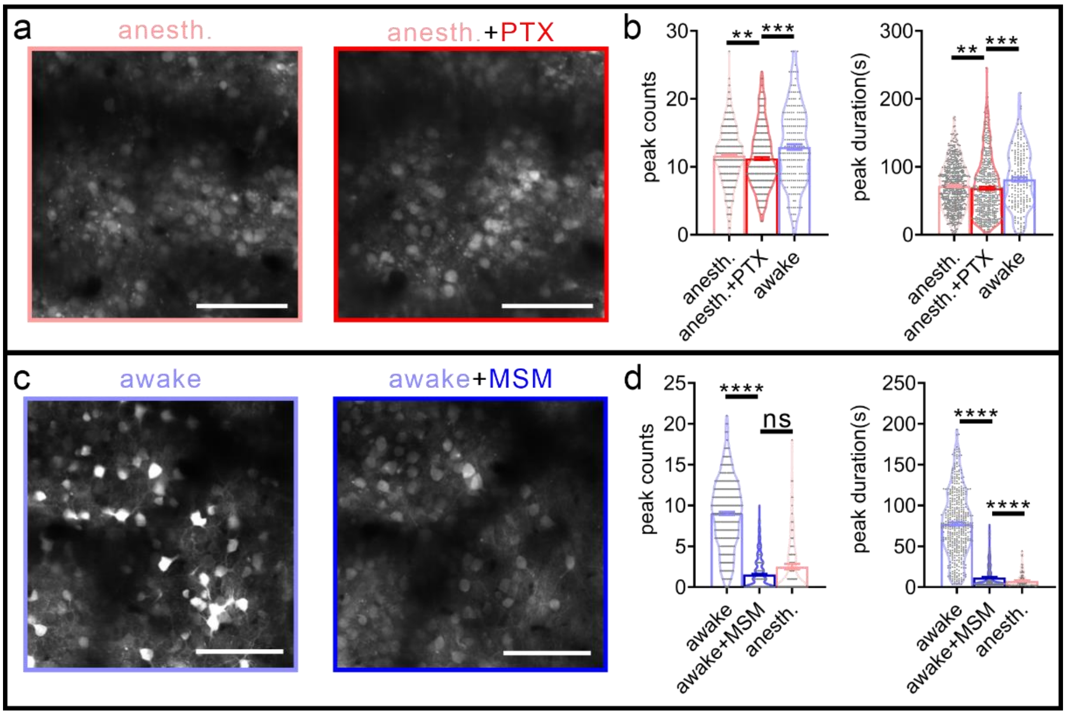


**Supplementary Fig. 6 Pharmacological manipulation alters spontaneous neuronal calcium activity.**

(a) Representative imaging field before and after picrotoxin (PTX) application. Scale bar, 100 μm. (b) Spontaneous calcium-activity measures in anesthetized mice before and after PTX, with awake measurements shown for reference. Peak count: 11.65 ± 0.17 (anesthetized) vs. 11.22 ± 0.21 (anesthetized + PTX), , p = 0.0073; 11.22 ± 0.21 (anesthetized + PTX) vs. 12.92 ± 0.42 (awake), p = 0.0003. Peak duration: 71.96 ± 1.37 s (anesthetized) vs. 68.63 ± 2.00 s (anesthetized + PTX), p = 0.0044; 68.63 ± 2.00 s (anesthetized + PTX) vs. 81.58 ± 3.14 s (awake), p = 0.0001. (c) Representative imaging field before and after muscimol (MSM) application. Scale bar, 100 μm. (d) MSM markedly reduces spontaneous calcium activity in awake mice. Peak count: 9.03 ± 0.20 (awake) vs. 1.55 ± 0.10 (awake + MSM), p < 0.0001; 1.55 ± 0.10 (awake + MSM) vs. 2.52 ± 0.34 (anesthetized), p = 0.1547. Peak duration: 77.56 ± 1.91 s (awake) vs. 11.76 ± 0.65 s (awake + MSM), p< 0.0001; 11.76 ± 0.65 s (awake + MSM) vs. 7.60 ± 0.84 s (anesthetized), p < 0.0001. Unpaired Mann-Whitney test. Data are presented as mean ± SEM. Related to Figs. 4 and 5.

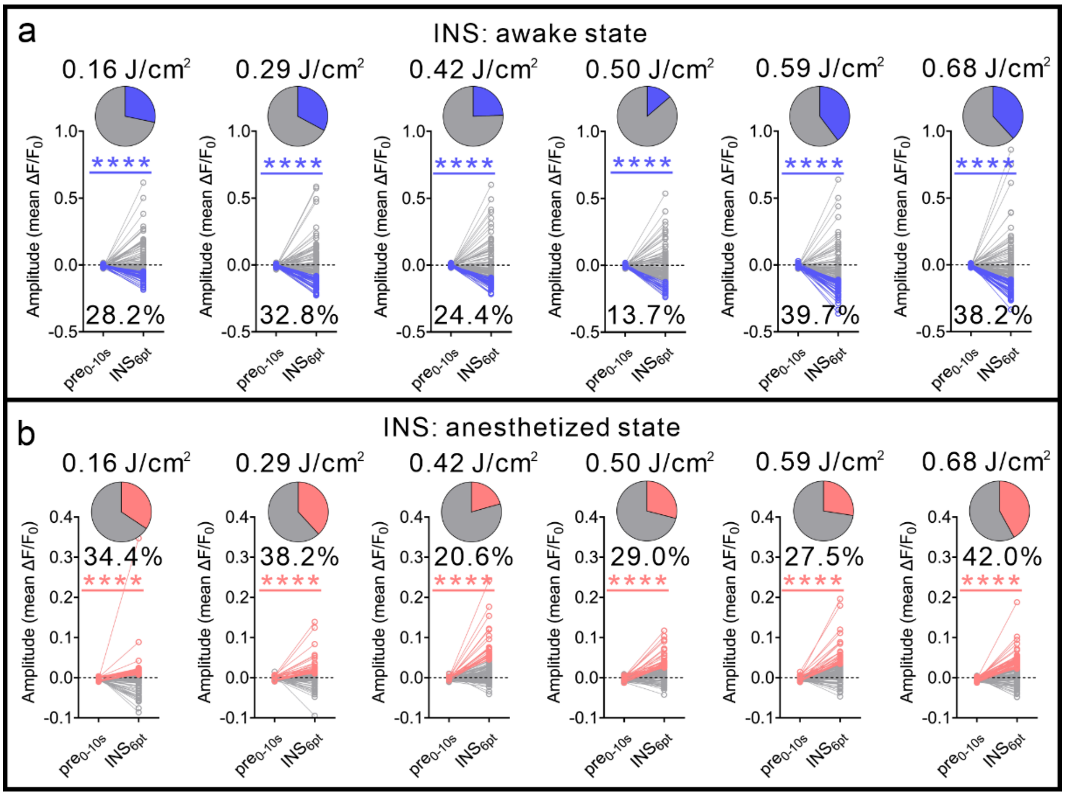


**Supplementary Fig. 7 Distinct neuronal response patterns to INS across varying stimulation intensities under awake and anesthetized conditions.**

(a) Neurons in the selected awake-response group exhibit negative INS-evoked responses across the tested stimulation intensities. (b) Neurons in the selected anesthetized-response group exhibit positive INS-evoked responses. Statistical analyses were performed using the paired Wilcoxon test. Data are presented as mean ± SEM and summarized in Supplementary Table 7. These groups are response-defined and should not be interpreted as biological neuronal subclasses. Related to Fig. 6.

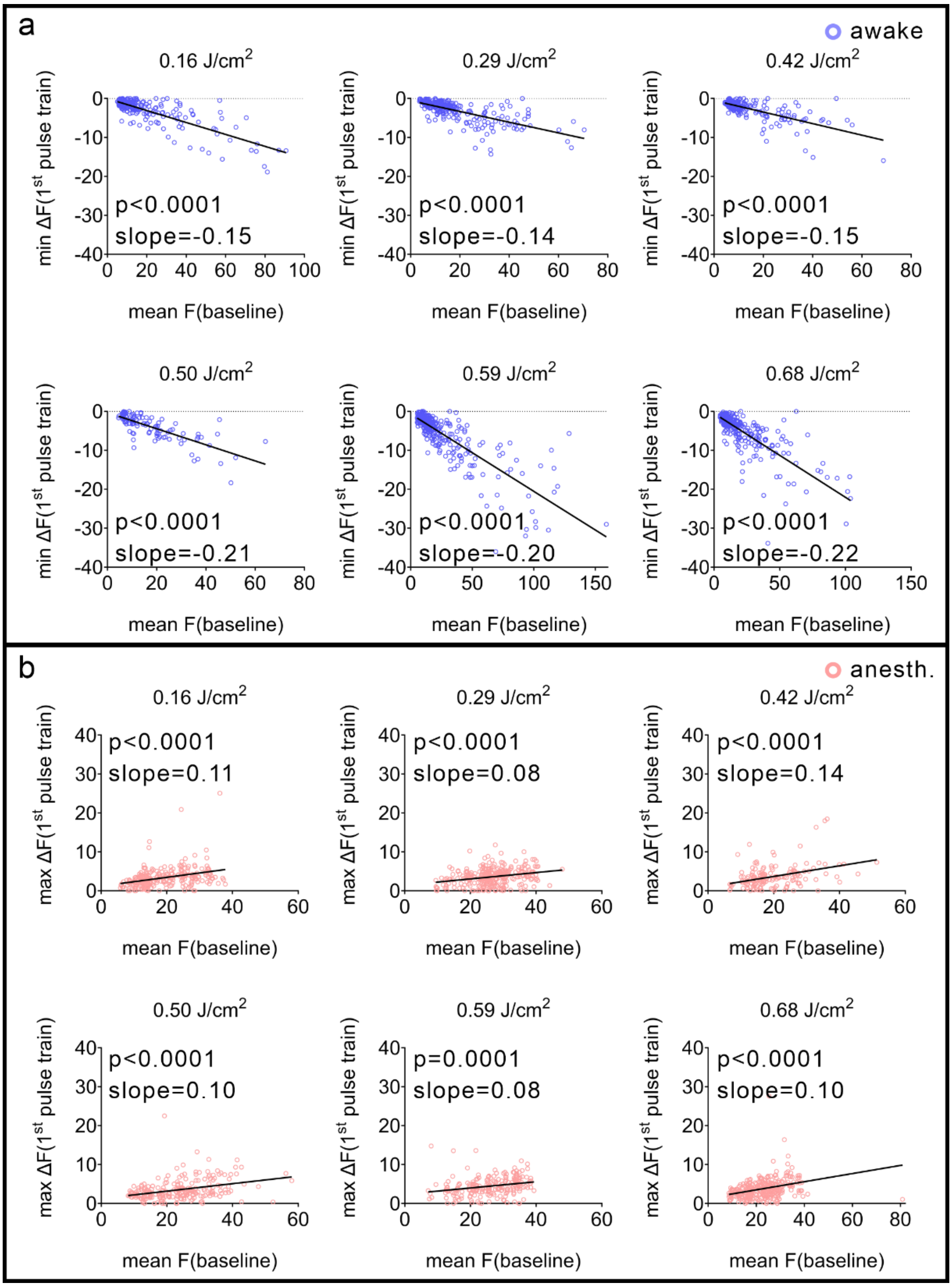


**Supplementary Fig. 8 Association between pre-INS fluorescence level and neuronal responses to the first INS train across stimulation intensities.**

(a) In the selected awake-response group, neuronal responses show negative associations with the pre-INS fluorescence level across the tested INS intensities (blue traces). (b) In the selected anesthetized-response group, positive associations are observed across the tested intensities (red traces). These regressions are exploratory because repeated trials are nested within neurons and animals. Linear-regression statistics are summarized in Supplementary Table 8. Related to Fig. 6.

**Supplementary Table 1. Correlation and amplitude values for neuronal responses to INS in awake and anesthetized states.**

| | Correlation | | Amplitude | |
|---|---|---|---|---|
| Intensity | awake | anesth. | awake | anesth. |
| Paired values (n = 131 neurons) | | | | |
| 0.16 J/cm$^2$ | -0.0021 ± 0.0089 | 0.0411 ± 0.0066 | 0.0104 ± 0.0104 | 0.00003 ± 0.0033 |
| 0.29 J/cm$^2$ | -0.0603 ± 0.0111 | 0.0674 ± 0.0095 | -0.0181 ± 0.0122 | 0.0038 ± 0.0027 |
| 0.42 J/cm$^2$ | -0.0433 ± 0.0124 | 0.1461 ± 0.0121 | -0.0146 ± 0.0121 | 0.0229 ± 0.0034 |
| 0.50 J/cm$^2$ | -0.0838 ± 0.0138 | 0.1328 ± 0.0141 | -0.0259 ± 0.0109 | 0.017 ± 0.0022 |
| 0.59 J/cm$^2$ | -0.071 ± 0.016 | 0.079 ± 0.0169 | -0.0622 ± 0.0126 | 0.0225 ± 0.003 |
| 0.68 J/cm$^2$ | -0.1015 ± 0.0175 | 0.1027 ± 0.0186 | -0.0407 ± 0.0152 | 0.0164 ± 0.0027 |
| 0.76 J/cm$^2$ | -0.1334 ± 0.0163 | 0.1393 ± 0.02 | -0.059 ± 0.0126 | 0.0246 ± 0.0029 |
| Comparison | INS vs. sp | INS vs. sp | INS vs. sp | INS vs. sp |
| Paired Wilcoxon test | | | | |
| 0.16 J/cm$^2$ | $p = 0.0016$ | $p = 0.0004$ | $p = 0.2717$ | $p = 0.0424$ |
| 0.29 J/cm$^2$ | $p < 0.0001$ | $p < 0.0001$ | $p = 0.0297$ | $p < 0.0001$ |
| 0.42 J/cm$^2$ | $p < 0.0001$ | $p < 0.0001$ | $p = 0.1486$ | $p < 0.0001$ |
| 0.50 J/cm$^2$ | $p < 0.0001$ | $p < 0.0001$ | $p = 0.0081$ | $p < 0.0001$ |
| 0.59 J/cm$^2$ | $p < 0.0001$ | $p = 0.0007$ | $p < 0.0001$ | $p < 0.0001$ |
| 0.68 J/cm$^2$ | $p < 0.0001$ | $p < 0.0001$ | $p = 0.0003$ | $p < 0.0001$ |
| 0.76 J/cm$^2$ | $p < 0.0001$ | $p < 0.0001$ | $p < 0.0001$ | $p < 0.0001$ |

**Supplementary Table 2. Correlation and amplitude values for neuronal responses to INS in Up and Other neuron groups.**

| | Correlation: Up neuron | |
|---|---|---|
| Intensity | awake | anesth. |
| 0.16 J/cm$^2$ | 0.0294 ± 0.0871, n = 10 | 0.0784 ± 0.0904, n = 10 |
| 0.29 J/cm$^2$ | -0.0584 ± 0.1621, n = 35 | 0.0441 ± 0.1139, n = 35 |
| 0.42 J/cm$^2$ | -0.0753 ± 0.1728, n = 30 | 0.1513 ± 0.1511, n = 30 |
| 0.50 J/cm$^2$ | -0.1309 ± 0.1978, n = 30 | 0.1038 ± 0.1634, n = 30 |
| 0.59 J/cm$^2$ | -0.1134 ± 0.2040, n = 58 | 0.0725 ± 0.2138, n = 58 |
| 0.68 J/cm$^2$ | -0.1756 ± 0.2349, n = 50 | 0.0965 ± 0.2260, n = 50 |
| 0.76 J/cm$^2$ | -0.1806 ± 0.2163, n = 53 | 0.1387 ± 0.2169, n = 53 |
| | Amplitude: Up neuron | |
| Intensity | awake | anesth. |
| 0.16 J/cm$^2$ | -0.1406 ± 0.0293, n = 10 | -0.0128 ± 0.0311, n = 10 |
| 0.29 J/cm$^2$ | -0.1417 ± 0.0377, n = 35 | 0.0057 ± 0.0377, n = 35 |
| 0.42 J/cm$^2$ | -0.1419 ± 0.0312, n = 30 | 0.0145 ± 0.0285, n = 30 |
| 0.50 J/cm$^2$ | -0.1523 ± 0.0427, n = 30 | 0.0097 ± 0.0205, n = 30 |
| 0.59 J/cm$^2$ | -0.1635 ± 0.0573, n = 58 | 0.0150 ± 0.0262, n = 58 |
| 0.68 J/cm$^2$ | -0.1683 ± 0.0539, n = 50 | 0.0096 ± 0.0295, n = 50 |
| 0.76 J/cm$^2$ | -0.1806 ± 0.0554, n = 53 | 0.0205 ± 0.0329, n = 53 |
| | Correlation: Other neuron | |
| Intensity | awake | anesth. |
| 0.16 J/cm$^2$ | -0.0023 ± 0.1038, n = 112 | 0.0391 ± 0.0743, n = 112 |
| 0.29 J/cm$^2$ | -0.0487 ± 0.1126, n = 82 | 0.0610 ± 0.0966, n = 82 |
| 0.42 J/cm$^2$ | -0.0076 ± 0.1254, n = 81 | 0.1274 ± 0.1327, n = 81 |
| 0.50 J/cm$^2$ | -0.0646 ± 0.1410, n = 89 | 0.1365 ± 0.1679, n = 89 |
| 0.59 J/cm$^2$ | -0.0159 ± 0.1513, n = 61 | 0.0697 ± 0.1723, n = 61 |
| 0.68 J/cm$^2$ | -0.0449 ± 0.1610, n = 67 | 0.0985 ± 0.2094, n = 67 |
| 0.76 J/cm$^2$ | -0.0938 ± 0.1541, n = 62 | 0.1051 ± 0.2294, n = 62 |

| Amplitude: Other neuron | | |
|---|---|---|
| Intensity | awake | anesth. |
| 0.16 J/cm$^2$ | 0.0107 ± 0.1099, n = 112 | 0.0001 ± 0.0398, n = 112 |
| 0.29 J/cm$^2$ | -0.0090 ± 0.0920, n = 82 | 0.0010 ± 0.0290, n = 82 |
| 0.42 J/cm$^2$ | -0.0267 ± 0.0838, n = 81 | 0.0239 ± 0.0385, n = 81 |
| 0.50 J/cm$^2$ | -0.0170 ± 0.0842, n = 89 | 0.0181 ± 0.0252, n = 89 |
| 0.59 J/cm$^2$ | -0.0280 ± 0.0783, n = 61 | 0.0253 ± 0.0352, n = 61 |
| 0.68 J/cm$^2$ | -0.0129 ± 0.1130, n = 67 | 0.0161 ± 0.0253, n = 67 |
| 0.76 J/cm$^2$ | -0.0303 ± 0.0485, n = 62 | 0.0223 ± 0.0302, n = 62 |

**Supplementary Table 3. Amplitude values for neuronal responses to INS in PTX-manipulated anesthetized mice.**

| Intensity | $Pre_{0\text{-}10s}$ | $INS_{6pt}$ | Paired Wilcoxon test |
|---|---|---|---|
| Amplitude of neuron responses to INS in PTX-manipulated mice | | | |
| 0.16 $J/cm^2$ | -0.0001 ± 0.0000 | -0.0171 ± 0.0012 | n = 173, $p < 0.0001$ |
| 0.42 $J/cm^2$ | 0.0007 ± 0.0001 | -0.0802 ± 0.0038 | n = 142, $p < 0.0001$ |
| 0.59 $J/cm^2$ | 0.0007 ± 0.0001 | -0.1055 ± 0.0033 | n = 191, $p < 0.0001$ |
| 0.76 $J/cm^2$ | 0.0000 ± 0.0001 | -0.0838 ± 0.0031 | n = 222, $p < 0.0001$ |

**Supplementary Table 4. Linear regression for neuronal responses to INS in PTX-manipulated mice.**

| Intensity | 0 | 0.16 $J/cm^2$ | 0.42 $J/cm^2$ | 0.59 $J/cm^2$ | 0.76 $J/cm^2$ |
|---|---|---|---|---|---|
| Amplitude ($INS_{6pt}$) | 0.0173 ± 0.0032 | -0.0171 ± 0.0012 | -0.0802 ± 0.0038 | -0.1055 ± 0.0033 | -0.0838 ± 0.0031 |
| Linear regression | | | | | |
| | $Y = -0.1526 \times X + 0.005$ | | | $R^2 = 0.84$, $p = 0.03$ | |

**Supplementary Table 5. Amplitude values for neuronal responses to INS in MSM-manipulated awake mice.**

| Intensity | $Pre_{0\text{-}10s}$ | $INS_{6pt}$ | Paired Wilcoxon test |
|---|---|---|---|
| Amplitude of neuron responses to INS in MSM-manipulated mice | | | |
| 0.16 $J/cm^2$ | 0.0013 ± 0.0000 | 0.0351 ± 0.0009 | n = 161, $p < 0.0001$ |
| 0.42 $J/cm^2$ | 0.0016 ± 0.0000 | 0.0282 ± 0.0008 | n = 197, $p < 0.0001$ |
| 0.59 $J/cm^2$ | 0.0013 ± 0.0000 | 0.0442 ± 0.0010 | n = 278, $p < 0.0001$ |
| 0.76 $J/cm^2$ | 0.0014 ± 0.0000 | 0.0549 ± 0.0012 | n = 234, $p < 0.0001$ |

**Supplementary Table 6. Linear regression for neuronal responses to INS in MSM-manipulated mice.**

| Intensity | 0 | 0.16 $J/cm^2$ | 0.42 $J/cm^2$ | 0.59 $J/cm^2$ | 0.76 $J/cm^2$ |
|---|---|---|---|---|---|
| Amplitude ($INS_{6pt}$) | -0.0005 ± 0.0006 | 0.0351 ± 0.0009 | 0.0282 ± 0.0008 | 0.0442 ± 0.0010 | 0.0549 ± 0.0012 |
| Linear regression | | | | | |
| | | $Y = 0.0595 \times X + 0.0094$ | | $R^2 = 0.77$, $p = 0.0497$ | |

**Supplementary Table 7. Amplitude values for distinct neuronal responses to INS between awake and anesthetized states.**

| Intensity | $Pre_{0\text{-}10s}$ | $INS_{6pt}$ | Paired Wilcoxon test |
|---|---|---|---|
| Neuronal responses to INS in awake state | | | |
| 0.16 J/cm$^2$ | -0.0083 ± 0.0013 | -0.0905 ± 0.0059 | n = 37, p < 0.0001 |
| 0.29 J/cm$^2$ | -0.0029 ± 0.0011 | -0.1318 ± 0.0061 | n = 43, p < 0.0001 |
| 0.42 J/cm$^2$ | -0.0052 ± 0.0014 | -0.1390 ± 0.0057 | n = 32, p < 0.0001 |
| 0.50 J/cm$^2$ | -0.0017 ± 0.0023 | -0.1799 ± 0.0077 | n = 18, p < 0.0001 |
| 0.59 J/cm$^2$ | -0.0057 ± 0.0014 | -0.1701 ± 0.0079 | n = 52, p < 0.0001 |
| 0.68 J/cm$^2$ | -0.0069 ± 0.0012 | -0.1683 ± 0.0076 | n = 50, p < 0.0001 |
| 0.76 J/cm$^2$ | -0.0091 ± 0.0016 | -0.2011 ± 0.0076 | n = 40, p < 0.0001 |
| Neuronal responses to INS in anesthetized state | | | |
| 0.16 J/cm$^2$ | -0.0013 ± 0.0006 | 0.0237 ± 0.0076 | n = 45, p < 0.0001 |
| 0.29 J/cm$^2$ | -0.0037 ± 0.0007 | 0.0259 ± 0.0036 | n = 50, p < 0.0001 |
| 0.42 J/cm$^2$ | -0.0008 ± 0.0006 | 0.0813 ± 0.0090 | n = 27, p < 0.0001 |
| 0.50 J/cm$^2$ | -0.0044 ± 0.0009 | 0.0446 ± 0.0038 | n = 38, p < 0.0001 |
| 0.59 J/cm$^2$ | -0.0040 ± 0.0011 | 0.0598 ± 0.0063 | n = 36, p < 0.0001 |
| 0.68 J/cm$^2$ | -0.0047 ± 0.0007 | 0.0428 ± 0.0036 | n = 55, p < 0.0001 |
| 0.76 J/cm$^2$ | -0.0024 ± 0.0008 | 0.0556 ± 0.0037 | n = 51, p < 0.0001 |

**Supplementary Table 8. Linear regression for neuronal responses to INS of trial-by-trial data.**

| Intensity | State | Equation | $R^2$ and P value |
|---|---|---|---|
| 0.16 J/cm$^2$ | awake | Y = -0.1533×X - 0.0276 | $R^2$ = 0.64, p < 0.0001 |
| | anesth. | Y = 0.1126×X + 1.198 | $R^2$ = 0.12, p < 0.0001 |
| 0.29 J/cm$^2$ | awake | Y = -0.1375×X - 0.5819 | $R^2$ = 0.48, p < 0.0001 |
| | anesth. | Y = 0.0815×X + 1.41 | $R^2$ = 0.08, p < 0.0001 |
| 0.42 J/cm$^2$ | awake | Y = -0.1483×X - 0.5107 | $R^2$ = 0.47, p < 0.0001 |
| | anesth. | Y = 0.1366×X + 0.9491 | $R^2$ = 0.14, p < 0.0001 |
| 0.50 J/cm$^2$ | awake | Y = -0.2067×X - 0.3109 | $R^2$ = 0.58, p < 0.0001 |
| | anesth. | Y = 0.0959×X + 1.245 | $R^2$ = 0.12, p < 0.0001 |
| 0.59 J/cm$^2$ | awake | Y = -0.1987×X - 0.6649 | $R^2$ = 0.67, p < 0.0001 |
| | anesth. | Y = 0.08×X + 2.375 | $R^2$ = 0.07, p = 0.0001 |
| 0.68 J/cm$^2$ | awake | Y = -0.2154×X - 0.5161 | $R^2$ = 0.54, p < 0.0001 |
| | anesth. | Y = 0.1046×X + 1.403 | $R^2$ = 0.07, p < 0.0001 |
| 0.76 J/cm$^2$ | awake | Y = -0.2882×X + 0.6076 | $R^2$ = 0.75, p < 0.0001 |
| | anesth. | Y = 0.1533×X + 0.3523 | $R^2$ = 0.28, p < 0.0001 |
